# TAPAS: an open-source software package for Translational Neuromodeling and Computational Psychiatry

**DOI:** 10.1101/2021.03.12.435091

**Authors:** Stefan Frässle, Eduardo A. Aponte, Saskia Bollmann, Kay H. Brodersen, Cao T. Do, Olivia K. Harrison, Samuel J. Harrison, Jakob Heinzle, Sandra Iglesias, Lars Kasper, Ekaterina I. Lomakina, Christoph Mathys, Matthias Müller-Schrader, Inês Pereira, Frederike H. Petzschner, Sudhir Raman, Dario Schöbi, Birte Toussaint, Lilian A. Weber, Yu Yao, Klaas E. Stephan

## Abstract

Psychiatry faces fundamental challenges with regard to mechanistically guided differential diagnosis, as well as prediction of clinical trajectories and treatment response of individual patients. This has motivated the genesis of two closely intertwined fields: (i) Translational Neuromodeling (TN), which develops “computational assays” for inferring patient-specific disease processes from neuroimaging, electrophysiological, and behavioral data; and (ii) Computational Psychiatry (CP), with the goal of incorporating computational assays into clinical decision making in everyday practice. In order to serve as objective and reliable tools for clinical routine, computational assays require end-to-end pipelines from raw data (input) to clinically useful information (output). While these are yet to be established in clinical practice, individual components of this general end-to-end pipeline are being developed and made openly available for community use.

In this paper, we present the **T**ranslational **A**lgorithms for **P**sychiatry-**A**dvancing **S**cience (TAPAS) software package, an open-source collection of building blocks for computational assays in psychiatry. Collectively, the tools in TAPAS presently cover several important aspects of the desired end-to-end pipeline, including: (i) tailored experimental designs and optimization of measurement strategy prior to data acquisition, (ii) quality control during data acquisition, and (iii) artifact correction, statistical inference, and clinical application after data acquisition. Here, we review the different tools within TAPAS and illustrate how these may help provide a deeper understanding of neural and cognitive mechanisms of disease, with the ultimate goal of establishing automatized pipelines for predictions about individual patients. We hope that the openly available tools in TAPAS will contribute to the further development of TN/CP and facilitate the translation of advances in computational neuroscience into clinically relevant computational assays.

## 1 INTRODUCTION

Contemporary psychiatry uses disease classifications that are almost entirely based on syndromes (i.e., patterns of symptoms and signs) as defined by the Diagnostic and Statistical Manual of Mental Disorders (DSM; American Psychiatric Association, 2013) or the International Classification of Diseases (ICD; World Health Organization, 1990). While these schemes are valuable in that they provide a stratification of mental illness that relates to the subjective phenomenology of patients, they are inherently limited as they do not rest on pathophysiological or aetiological concepts of diseases. As a consequence, clinical labels proposed by DSM or ICD (e.g., schizophrenia or depression) typically have limited predictive validity with regard to clinical trajectories and do not inform the optimal treatment selection in individual patients (Cuthbert and Insel, 2010; Kapur et al., 2012). Furthermore, clinical and scientific evidence suggests that these labels do not describe distinct categorical entities, but rather spectrum disorders that are characterized by substantial heterogeneity and overlap (Krystal and State, 2014; Owen, 2014; Stephan et al., 2016a).

This has motivated novel approaches to advance our understanding of the pathophysiological and psychopathological processes underlying diseases, and to ultimately inform differential diagnosis and treatment prediction in individual patients (Stephan et al., 2016b). In addition to the rise of (epi)genetic approaches, advances in computational neuroscience have fueled hopes that it may become possible to establish quantitative diagnostic and prognostic computational tools that significantly improve clinical practice in psychiatry. In particular, mathematical models of neuroimaging data, as obtained using functional magnetic resonance imaging (fMRI) and electro/magnetoencephalography (EEG/MEG), hold great promise as they might offer readouts of the symptom-producing physiological processes underlying brain disorders (Adams et al., 2020; Deco and Kringelbach, 2014; Frässle et al., 2018b; Gilbert et al., 2016; Moran et al., 2011; Symmonds et al., 2018). Similarly, advances in computational models of human behavior may enable inference on psychopathological processes at the computational (information-processing) level (Cole et al., 2020; Iglesias et al., 2016; Lawson et al., 2017; Maia and Frank, 2011; Powers et al., 2017; Stephan and Mathys, 2014).

Efforts to exploit these scientific advances can be grouped into two separate yet overlapping approaches. Primarily methodological efforts towards the development of “computational assays” for inferring brain disease processes from neuroimaging, electrophysiological, and behavioral data are referred to as Translational Neuromodeling (TN); by contrast, Computational Psychiatry (CP), Computational Neurology (CN), and Computational Psychosomatics (CPS) are concerned with concrete applications in the respective clinical domains, with the ultimate goal of incorporating computational assays into routine clinical decision-making (Figure 1). Although many of the tools in TAPAS are equally useful for CN and CPS, here we focus on CP as this is arguably the most developed of the computational clinical neurosciences (Adams et al., 2016; Browning et al., 2020; Friston et al., 2014; Huys et al., 2016; Jirsa et al., 2016; Jirsa et al., 2010; Montague et al., 2012; Paulus et al., 2016; Stephan and Mathys, 2014; Stephan et al., 2015; Stephan et al., 2017; Wang and Krystal, 2014).

**Figure 1.**
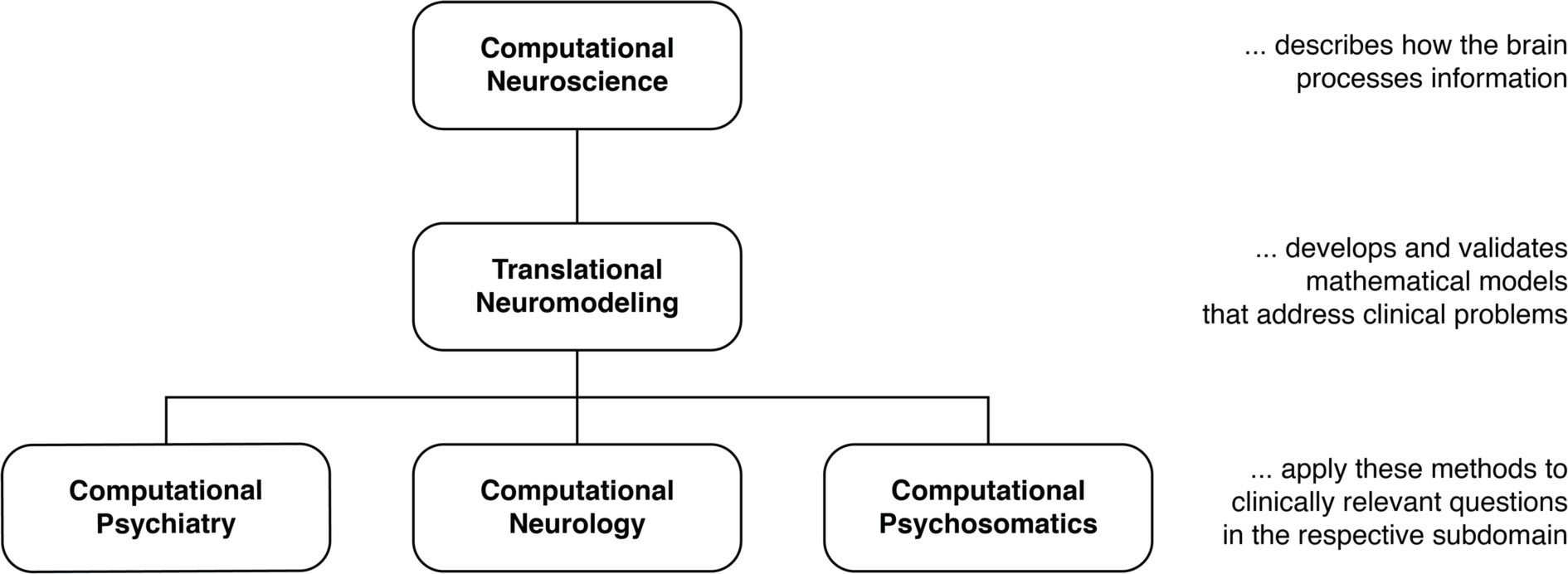
Taxonomy for different disciplines in the computational neurosciences and their relation to clinical questions. Translational Neuromodeling (TN) develops and validates mathematical models for addressing clinical problems, whereas Computational Psychiatry (CP), Neurology (CN), and Psychosomatics (CPS) then apply these methods to clinically relevant questions. Reprinted with permission from (Frässle et al., 2018b). Copyright 2018 Wiley.

Developments of computational assays are often based on generative models of neuroimaging or behavioral data. Generative models describe how measured data may have been caused by a particular (neuronal or cognitive) mechanism; their inversion allows computational assays to operate on inferred states of neural or cognitive systems (Browning et al., 2020; Stephan and Mathys, 2014). This mechanistic interpretability is crucial in many clinical contexts. Additionally, traditional machine learning (ML) plays a central role in TN/CP, for example by translating the inferences from computational assays into patient-specific predictions, an approach referred to as “generative embedding” (Brodersen et al., 2011).^1^

While early TN/CP proposals date back over a decade (e.g., see Stephan et al., 2006), computational assays are yet to enter into routine clinical practice in psychiatry. In order to achieve translational success, computational assays will have to build on automatized and validated end-to-end pipelines and tools for optimal data acquisition. These pipelines need to support a complete analysis stream that takes raw data as input, and outputs clinically actionable results that are derived from inferred latent (hidden) computational quantities with pathophysiological or psychopathological relevance. Such an end-to-end pipeline will incorporate a series of fundamental steps (Figure 2): (i) Design, (ii) Conduct, (iii) Check & Correct, (iv) Preprocessing, (v) Inference, (vi) Clinical application. Each of these components poses significant challenges given the complex nature of the acquired data and of the computational tools.

**Figure 2.**
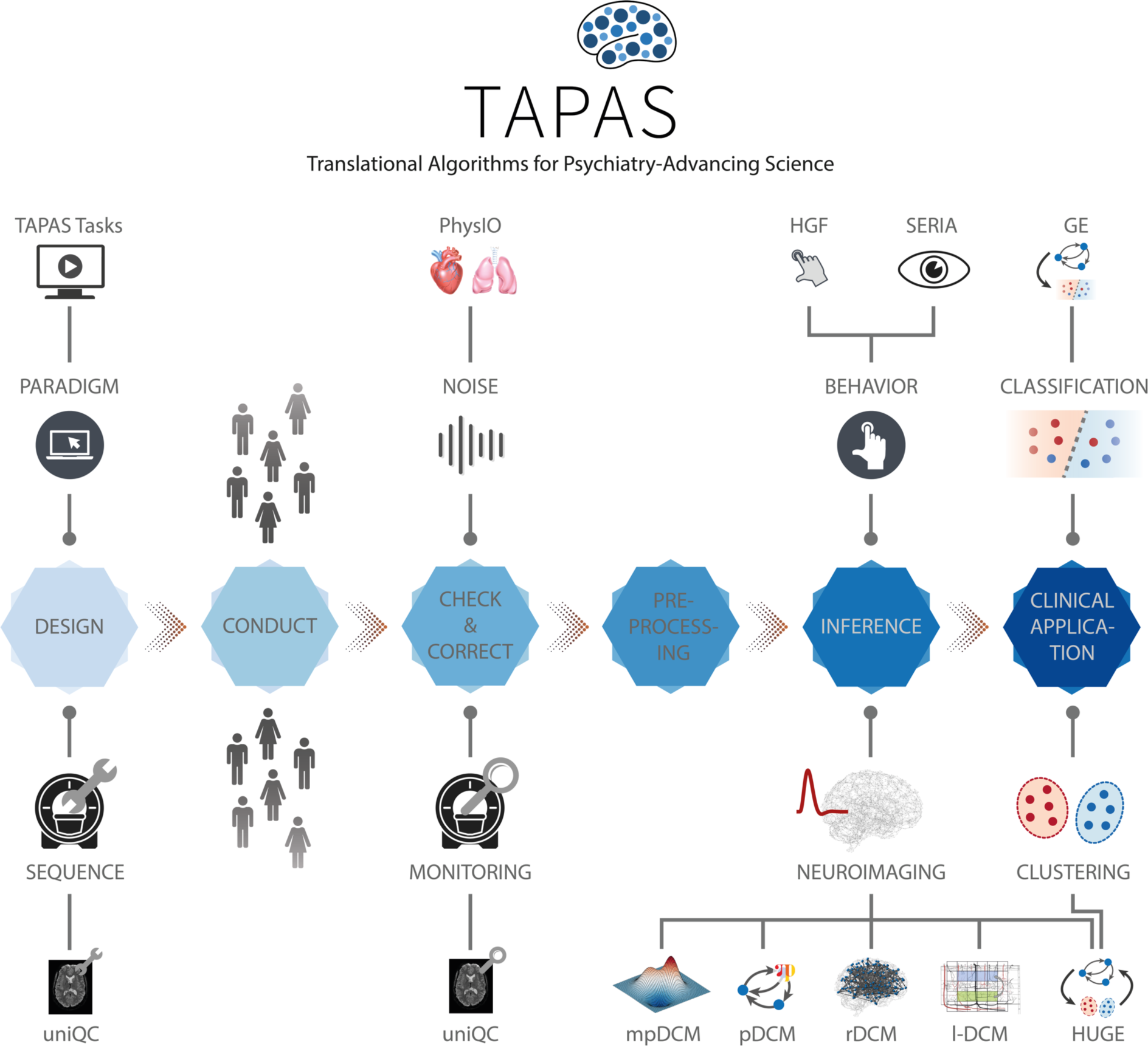
TAPAS components in a proposed end-to-end pipeline of a clinically relevant computational assay. This end-to-end pipeline will need to incorporate all steps from the raw imaging or behavioral data to a final clinical recommendation. Such a computational assays will capture at least the following crucial steps: (i) Design, (ii) Conduct, (iii) Check & Correct, (iv) Preprocessing, (v) Inference, and (vi) Clinical application. Various components of TAPAS feature into one or several of these steps and aim to address important questions and limitations that have so far hampered translational success.

Here, we review the current state of development towards such an end-to-end pipeline, with a particular focus on our own work and software.

Considerable efforts have recently been made to develop standardized and user-friendly software packages that could serve as individual components for computational assays. For instance, in the context of neuroimaging data, packages like Statistical Parametric Mapping (SPM; Friston et al., 2006), FMRIB Software Library (FSL; Smith et al., 2004), or Analysis of Functional NeuroImages (AFNI; Cox, 1996) are widespread tools and cover several aspects of the aforementioned end-to-end pipeline, including preprocessing of neuroimaging data and statistical inference (typically in the framework of General Linear Models). Similarly, for behavioral data, software packages like the VBA Toolbox (Daunizeau et al., 2014), hBayesDM (Ahn et al., 2017), KFAS (Helske, 2017), COMPASS (Yousefi et al., 2018), or HDDM (Wiecki et al., 2013) allow inference on computational (information processing) quantities. This list is non-exhaustive and more software packages could be mentioned. While all of these packages have proven highly valuable to study behavior and brain function in humans, none of them has been designed with the specific goal of constructing a pipeline for clinically useful computational assays.

In this paper, we focus on the **T**ranslational **A**lgorithms for **P**sychiatry-**A**dvancing **S**cience (TAPAS) software package which represents a collection of toolboxes that, collectively, aim to advance computational modeling of neuroimaging and behavioral data. While applicable to study human behavior and brain function in health, TAPAS differs from the aforementioned software packages in that its designated purpose is to provide clinically useful tools at every stage of the aforementioned end-to-end pipeline in order to advance translational success of computational approaches to psychiatry. TAPAS is primarily written in MATLAB (with some components in C and Python) and distributed as open-source code under the GNU General Public License 3.0 (https://www.translationalneuromodeling.org/tapas). It does not represent a single uniform toolbox but rather a collection of toolboxes, each of which addresses a specific problem that arises in TN/CP approaches to neuroimaging and/or behavioral data analysis (Figure 2). In brief, TAPAS contains: (i) tailored experimental paradigms (tasks) that probe psychopathologically and/or pathophysiologically relevant processes, (ii) tools for optimization and monitoring of data quality in the specific context of fMRI, (iii) model-based physiological noise correction techniques for fMRI data, and (iv) generative models and associated statistical techniques that enable inference on latent (hidden) neurophysiological or cognitive quantities from neuroimaging or behavioral data. The latter range from network/circuit models that infer effective (directed) connectivity from fMRI and EEG/MEG data to behavioral models that extract computational quantities from observed actions (e.g., decisions or eye movements). To facilitate usability of our software, TAPAS is complemented with comprehensive documentation for each toolbox, as well as an active forum where users can seek help (https://github.com/translationalneuromodeling/tapas/issues).

Here, we provide a general overview of the different software toolboxes included in TAPAS and highlight how these may support the development of clinically useful computational assays for psychiatry. The paper is not meant to provide a comprehensive description of each toolbox, but instead offers a high-level perspective on how the different tools relate to each other in order to jointly advance TN/CP. For readers interested in a more in-depth treatment of a particular toolbox, references will be provided in the respective sections.

## 2 DESIGN

The development of carefully designed experimental manipulations and the acquisition of high-quality data is paramount for (clinical) modeling. This is because any conclusion – whether a scientific or clinical one – fundamentally rests on the underlying data. The goal of tailored experimental paradigms and optimized data acquisition is to increase both the sensitivity and specificity of clinical tests; this necessitates optimizations at different stages of data collection.

*(I) Prior to data collection*: Tailored experimental paradigms have to be designed that capture relevant processes of interest. This may relate to physiological and cognitive aspects in health, or to pathophysiological and psychopathological mechanisms in disease. Furthermore, optimization of the data acquisition process is vital to ensure high quality measurements. This is particularly important in the context of fMRI data, where it is common that project-specific MR sequences have to be developed. These aspects will be covered in the current section.
*(II) During data collection*: Measures have to be taken that allow maintaining a consistently high level of data quality across acquisitions; for instance, across different patients, scanners or sites. This is vital in order to ensure that comparable (clinical) conclusions can be drawn from the data. Tools that address this aspect of data quality control will be covered in section 0.
*(III) After data collection*: Post-hoc assessment of data quality is important to identify datasets that need to be excluded or extra analysis steps to deal with artifacts. Poor data quality might be due to severe artifacts and/or high noise levels in the data, induced by both the MR system and the participant itself (motion, physiological noise). To identify such cases, tools are required that allow for quantitative assessment of data quality and that facilitate the decision process as to whether satisfactory data quality can be restored or not. Finally, user-friendly tools are needed that enable automatized corrections to clean-up data as best as possible. We will address these aspects in section 3.

All these efforts ideally interact seamlessly with each other in order to maximize the sensitivity and specificity of diagnostic/prognostic tests that build on the acquired data. Suboptimal data acquisition and quality control can result in a high proportion of datasets that have to be excluded from further analyses or lead to false conclusions, which is particularly problematic in the context of clinical applications.

### 2.1 Harmonization of experimental design

For probing disease-relevant cognitive processes, a plethora of experimental tasks have been proposed that frequently only differ in small details. While some variations are valuable as they address somewhat different aspects of a cognitive process, this diversity complicates exact comparison of findings across studies and often gives rise to “approximate replications” where an initial finding is not replicated exactly but some (vaguely) related finding is linked to the original observation (Kapur et al., 2012; Maxwell, 2004). A prominent example of this in the context of clinical neuroimaging is the frontal dysfunction hypothesis in schizophrenia. Since the original report (Carter et al., 1998), several studies have re-examined this question using somewhat different experimental approaches and have reported a variety of different outcomes, ranging from hyper-to hypoactivation, to no obvious alterations at all (cf. Kapur et al., 2012; Van Snellenberg et al., 2006). While it is possible that these differences could be of pathophysiological relevance, potentially referring to different subgroups in the schizophrenic spectrum, inconsistencies in the utilized experimental manipulations render any differences difficult to interpret. Hence, until such variations are properly explained or controlled for, approximate replications do not provide a solid basis for clinical tests.

One way to address this challenge is by openly sharing established experimental tasks. Notably, while the call for open sharing of data has been very prominent in recent years (Poldrack and Gorgolewski, 2014; Poline et al., 2012; Van Horn et al., 2001), this is less the case for sharing the experimental tasks themselves (but see, for instance, the task protocols utilized by the Human Connectome Project (Van Essen et al., 2013) which have been shared openly).

To this end, TAPAS comprises the module “TAPAS Tasks” which represents a collection of experimental paradigms that have been designed and thoroughly tested (Figure 3, *top left*). TAPAS Tasks comprises several tasks that cover both the exteroceptive and interoceptive^2^ domains (for a complete list, see Table 1). For all paradigms, the stimulus code is provided as well as detailed documentation. This includes a comprehensive description of (i) the experimental task, (ii) software requirements, (iii) experimental set-up (including a list of necessary peripheral devices), and (iv) information on how to run the task.

**Figure 3.**
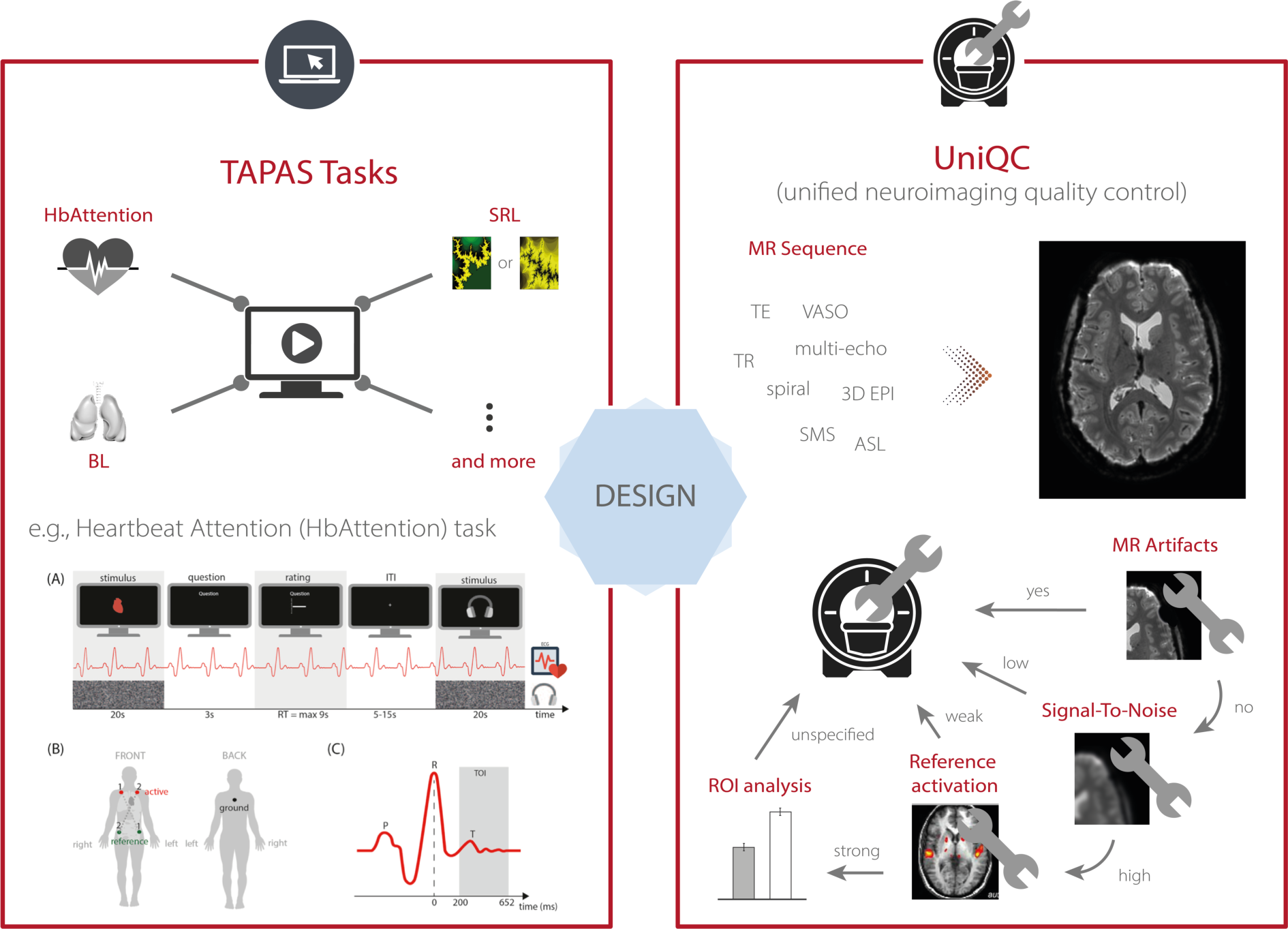
TAPAS components that aim to enhance data quality for scientific and clinical applications. This includes (*left, top*) TAPAS Tasks, a collection of experimental paradigms that have been devised and carefully tested, for instance, the Heartbeat Attention (HbAttention), stimulus-reward learning (SRL), and breathing learning (BL) task. For a complete list of tasks that are already included in TAPAS Tasks or will be included in one of the upcoming releases, see Table 1. (*Left, bottom*) Schematic overview of the experimental paradigm of the HbAttention task, as well as the placement of ECG electrodes and a typical ECG signal associated with a heartbeat (reprinted with permission from Petzschner et al., 2019). (*Right*) Furthermore, TAPAS comprises the unified neuroimaging quality control (UniQC) toolbox which is designed to facilitate the development and optimization of MR acquisition sequences. UniQC can assess (compute & visualize) different image quality metrics (IQMs) at different stages of an fMRI experiment (i.e., from raw data to statistical images). This facilitates the acquisition of high-quality data by implementing an iterative optimization process, including basic artifact checks, temporal stability analysis, functional sensitivity analyses in the whole brain or in particular regions of interest. UniQC enables this optimization in a highly flexible fashion, independent of the exact input data (e.g., sequence, dimensionality).

**Table 1.** List of tasks included in TAPAS Tasks. TAPAS Tasks includes a variety of different paradigms that probe exteroceptive and/or interoceptive processes. Note that some of the tasks listed here are not yet available in TAPAS, but will be included once the associated paper has been published.

| Task | Description |
| --- | --- |
| Heartbeat Attention (*HbAttention*) | The HbAttention task probes differences in neural responses to heartbeats due to changes in attentional focus. The paradigm consists of alternating conditions where participants focus attention either on their heart (interoceptive condition) or on an external sound stimulus (exteroceptive condition), while keeping the sensory stimulation identical (Petzschner et al., 2019). The HbAttention task requires the acquisition of an ECG and can be run in the context of an EEG experiment, measuring a Heartbeat Evoked Potential (HEP), or an fMRI experiment. |

<sup>2</sup> Exteroception refers to perception of sensations originating from the external world, whereas interoception refers to perception of sensations originating from the own body or “internal world”.

|  |  |
| --- | --- |
|  | The task is programmed such that the stimulus timing can be easily adjusted to the experimental modality. |
| Heartbeat Feedback (*HbFeedback*) | The HbFeedback task presents auditory-visual stimuli that are either locked to an individual's online detected heartbeat (veridical feedback about the heartbeat) or presented at a rate that is faster or slower than the individual's heartrate. The task assesses the effects of veridical versus false feedback on physiological and neural signals related to heartbeats. It requires the simultaneous recording of EEG and ECG signals. |
| Heartbeat Mismatch (*HbMMN*) | The HbMMN task consists of an auditory omission paradigm where a 'standard' tone is presented shortly after each heartbeat, but occasionally omitted ('deviant'). In different conditions, the delay between heartbeat and tone is varied. This allows to measure changes in stimulus-evoked and heartbeat-evoked potentials between standards and omissions. In a control condition, tone presentation times are unrelated to heartbeats. EEG and ECG signals are simultaneously recorded during the task. |
| Filter detection (*FD*) | The FD task is a perceptual threshold breathing task where participants have to indicate on each trial whether a very small resistance (i.e., filter) or sham (i.e., empty filter) was applied to the breathing system (yes/no version) (Harrison et al., 2020a), or in which interval resistance was applied (two-interval forced choice version). The task is tailored to assessing respiratory interoceptive accuracy and metacognition in individual participants. Behavioral responses are recorded. |
| Breathing learning (*BL*) | The BL task represents an associative learning task where participants learn the association between visual cues and the subsequent presence/absence of an inspiratory resistive load. Respiratory load is applied using a novel MRI-compatible breathing system that allows for remote administration and monitoring of resistive loads and whose construction plan has been published (Rieger et al., 2020). fMRI signals are recorded during the task. |
| Stimulus-reward learning (*SRL*) | The SRL task requires participants to predict which of two simultaneously presented visual stimuli (i.e., fractals) would yield a monetary reward. The association strengths between the visual cues and monetary outcomes change over the course of the experiment, introducing volatility. fMRI or EEG data can be acquired during the task. |
| Auditory Mismatch Negativity (aMMN) | The aMMN task is a variant of the auditory oddball paradigm in which the degree of volatility in the auditory stream varies over time. While engaging in a visual distraction task, participants passively listen to repeated presentations of a high and a low tone. During stable phases of the experiment, one stimulus reliably serves as the 'standard' (more frequent) tone and the other one as the 'deviant' tone. During volatile phases, the roles of standard and deviant switch more rapidly. Deviance processing can be compared between different levels of |
|  | stability/volatility (Weber et al., *in preparation*). Task versions are available for both EEG or fMRI recordings. |
| Visual Mismatch Negativity (vMMN) | The vMMN task implements the identical probabilistic stimulus sequence as the aMMN task (Do et al., *in preparation*). However, instead of auditory stimuli, Gabor patches of different orientations are used to probe mismatch responses in the visual domain. Task versions are available for both EEG and fMRI recordings. |
| Antisaccades (AS) | The AS task asks participants to perform antisaccades which are a type of voluntarily controlled eye movements. In TAPAS, code is available to run two different versions of the AS task (Aponte et al., 2019; 2018) using an EyeLink (SR Research, Ottawa, ON, Canada) eye tracking system. The versions of the task differ in the timing (with or without a delay before the eye movement) and position (at center or at peripheral target position) of the presentation of the task cue. |

Here, as an example, we describe the *Heartbeat Attention (HbAttention)* task in more detail (Figure 3, *bottom left*). The task implements a novel paradigm to probe purely attentional differences of the heartbeat evoked potential between exteroceptive and interoceptive conditions (Petzschner et al., 2019). The paradigm consists of alternating conditions where participants are asked to focus attention either on their heart or on a simultaneously presented auditory stimulus (white noise). Importantly, in both conditions the sensory stimulation is identical. Using this paradigm, Petzschner et al. (2019) found an increased heartbeat evoked potential during interoceptive compared with exteroceptive attention. A non-invasive readout of the attentional modulation of interoceptive processes could potentially be of high clinical relevance, since alterations in interoceptive processing have been recognized as a major component of various psychiatric conditions, including mood and anxiety disorders, eating disorders, drug addiction, as well as depression (Khalsa et al., 2018).

TAPAS Tasks represents work in progress, and newly devised experimental paradigms will be added to the module on a regular basis. We hope that by making these tasks openly available to the community, TAPAS Tasks may contribute to growing a collection of standardized experimental paradigms in TN/CP.

### 2.2 Optimization of MRI protocols

In the context of neuroimaging, data quality also depends heavily on the MR scanner settings and acquisition sequence. Carefully crafted sequences with optimized parameter choices can offer considerable gains in functional sensitivity and specificity for the targeted research question (Bollmann and Barth, 2020; Poser and Setsompop, 2018; Weldon and Olman, 2021). However, due to the large variety of available parameters and their interdependency, optimization of MR protocols is challenging and suboptimal acquisition choices might reduce data quality, e.g., low signal-to-noise ratio or pronounced artifacts like ghosting, ringing, signal dropouts and distortions due to magnetic field inhomogeneities (for an overview, see Bellon et al., 1986; Bright and Murphy, 2017; Kirilina et al., 2016; Volz et al., 2019). Hence, the development, optimization and validation of robust and powerful MR protocols prior to data acquisition is critical and tools are needed that ease this process.

To this end, TAPAS includes the *unified neuroimaging quality control* (UniQC) toolbox (Kasper*, Bollmann* et al., *in preparation*), which provides a framework for flexible, interactive and user-friendly computation and visualization of various quality measures in neuroimaging data (Figure 3, *right*). UniQC facilitates fast prototyping and optimization of acquisition sequences by providing tools for artifact detection and sensitivity analyses across the entire image or tailored towards specific regions of interest.

As sequence development constitutes an iterative process, feedback on image quality has to be immediate and specific to the protocol changes, so that the performed quality control (QC) query informs the operator on how to adjust parameters for the next scan (Figure 3, *right*). For example, if an unexpected bias field occurs in the mean image, both excitation and receiver channels could be compromised. In this case, fast display of individual coil images is critical, which would be omitted if the mean image were inconspicuous. Thus, QC during sequence development resembles a decision tree, where the outcome of one image quality metric (IQM) determines the selection of the next one, with varying display options. This necessitates an interactive, fast and flexible way to compute and visualize IQMs. Typically, such a decision tree starts from basic artifact checks over temporal stability all the way to functional sensitivity analyses in particular regions of interest, with the occasional return to the scanner, if a QC step fails (Figure 3, *right*).

To achieve this functionality, UniQC exploits the object-oriented programming capabilities in MATLAB. Importantly, UniQC is not restricted to 4-dimensional neuroimaging data (i.e., space and time) like most other software packages, but generalizes operations to n-dimensional data. This generalization to arbitrary numbers of dimensions comes in useful when handling data from multiple receiver coils (Hesselmann et al., 2004), as well as multi-echo (Poser et al., 2006; Posse et al., 1999) or combined magnitude/phase fMRI data (Kundu et al., 2017; Menon, 2002; Rowe, 2005) in a single unified framework. Similarly, prominent non-BOLD fMRI contrasts rely on an additional tag/control dimension, for instance, Vascular Space Occupancy (VASO; Huber et al., 2017; Lu et al., 2013) or Arterial Spin Labeling (ASL; Alsop et al., 2015; Koretsky, 2012), with the former being particularly relevant for depth-dependent fMRI. Furthermore, UniQC allows seamless integration with SPM and other MATLAB toolboxes in order to benefit from image and fMRI processing algorithms that are already available in those packages.

In summary, UniQC provides a flexible, interactive and user-friendly toolbox for evaluating MR pulse sequence development and quality control of n-dimensional neuroimaging data. These efforts carry over from the design stage to the data collection, in that UniQC allows utilizing the processing and visualization pipeline established here directly as a quality control protocol – enabling unique QC towards the aims of each study.

## 3 CONDUCT, CHECK AND CORRECT

Besides optimization steps prior to data acquisition, further steps are necessary during and after the measurement to ensure adequate data quality. Specifically, ongoing monitoring of data quality during image acquisition is critical, because both the MR system and the study participant constitute significant potential noise sources. On the system side, data quality across different time points and scanning sites may vary due to potential malfunctions or alterations in the scanner hardware, that must be detected in a timely manner. On the participant’s side, even under ideal circumstances, with tailored experimental designs, optimized acquisition sequences and thorough quality control measures, fMRI data is still subject to artifacts outside the experimenter’s control (e.g., motion, physiology; Glover et al., 2000; Kruger and Glover, 2001). Adequately correcting for these artifacts is essential to avoid bias in subsequent data analyses and to ensure that conclusions are not confounded (e.g., Power et al., 2012; Reuter et al., 2015; Yendiki et al., 2014). In what follows, we elaborate on these points and describe tools in TAPAS that address these challenges.

### 3.1 Quality monitoring

fMRI analyses rely on the content of brain (or spinal cord) images as information source. Thus, the visual inspection of raw images by one or multiple experts is still often considered a gold standard for quality control. However, visual assessment depends on individual rater experience and visualization choices (e.g., slice orientation, windowing) which may reduce the apparent information content of an image to detect artifacts or improper acquisition parameters (Gardner et al., 1995) and generally aggravates inter-rater reliability. Furthermore, limited time resources and fatigue make the naïve visual inspection of every raw image quickly unfeasible, as even a single fMRI dataset contains hundreds of volumes with dozens of slices each. This challenge is exacerbated for large-scale datasets, like the Human Connectome Project (HCP; Van Essen et al., 2013) or UK Biobank (Sudlow et al., 2015), where thousands of participants are measured.

Automation of quality control is therefore required, and can in principle target both aspects of the manual image classification by raters. A first approach is to replace raters by a machine learning algorithm working on derived IQMs to reduce the high-dimensional feature space of image time series. A second option is that the expertise of the rater can be harnessed more efficiently by providing flexible tools for image manipulation and visualization with intuitive interfaces to derive and inspect relevant IQMs, thereby reducing operator fatigue and inconsistency.

The first approach was explored early on for anatomical T1-weighted images (Woodard and Carley-Spencer, 2006) and demonstrated good discriminability of undistorted, noisy, and distorted images based on a subset of 239 IQMs. Since then, various additional tools for automatic quality control have been proposed, introducing additional IQMs to assess image quality (Gedamu et al., 2008; Mortamet et al., 2009; Pizarro et al., 2016). These methods have been refined and extended into scalable QC frameworks for large-scale fMRI studies, most notably in the form of the MRI Quality Control tool (MRIQC; Esteban et al., 2017) and within the UK biobank study (Alfaro-Almagro et al., 2018). Their key advancement lies in the ability to classify images in a binary fashion (“good” vs “problematic”) or even categorizing multiple artifact classes (Alfaro-Almagro et al., 2018). Standardized quality reports then provide guidance, as to whether data from a given participant should be included in subsequent analyses or not. In order to ensure accurate classification, these algorithms are typically trained on large curated datasets that include both patients and healthy controls (e.g., ABIDE (Di Martino et al., 2014) and DS030 (Poldrack et al., 2016) for MRIQC).

While such fully automatized approaches with minimal visual output and manual assessment might be the only viable solution in studies with thousands of participants, the required high degree of standardization of the acquisition protocol as well as the need for large, representative training datasets poses limitations on its utility. In TN/CP, less well-studied clinical populations and novel technologies can pose challenges when trying to exploit established mappings between IQMs and image quality. In particular, this can occur when employing advanced imaging hardware or acquisition sequences to maximize sensitivity for individual subject measurements, IQMs may fall outside the standard range or become inapplicable. For example, higher magnetic field strengths and customized high-density or surface coils (Hendriks et al., 2020; Keil and Wald, 2013) induce atypical image intensity variations (bias fields). Similarly, Nyquist ghosts manifest differently for spiral readouts than in conventional Cartesian echo-planar imaging. Thus, fMRI data may require different IQMs or thresholds when deviating from standard acquisition choices.

In these domains, the second option for QC automation is preferable. This approach empowers the rater to determine which IQMs to inspect, how to visualize them, and at which stage of the analysis stream to assess them. While parts of this approach have been implemented by providing standardized visual reports (e.g., MRIQC, fMRWhy) or interactive QC visualization tools (e.g., visualQC), a comprehensive framework that integrates all these functionalities has been missing. To address this, we designed the UniQC toolbox to meet these demands by offering flexible, interactive and user-friendly assessment of fMRI data (Figure 4, *left*). Importantly, the quality control pipeline derived during the sequence design stage (section 2) can be readily deployed to cover QC automation, and once quality issues are detected, UniQC also provides a framework to ‘interrogate’ the data efficiently and identify potential causes of the problem.

**Figure 4.**
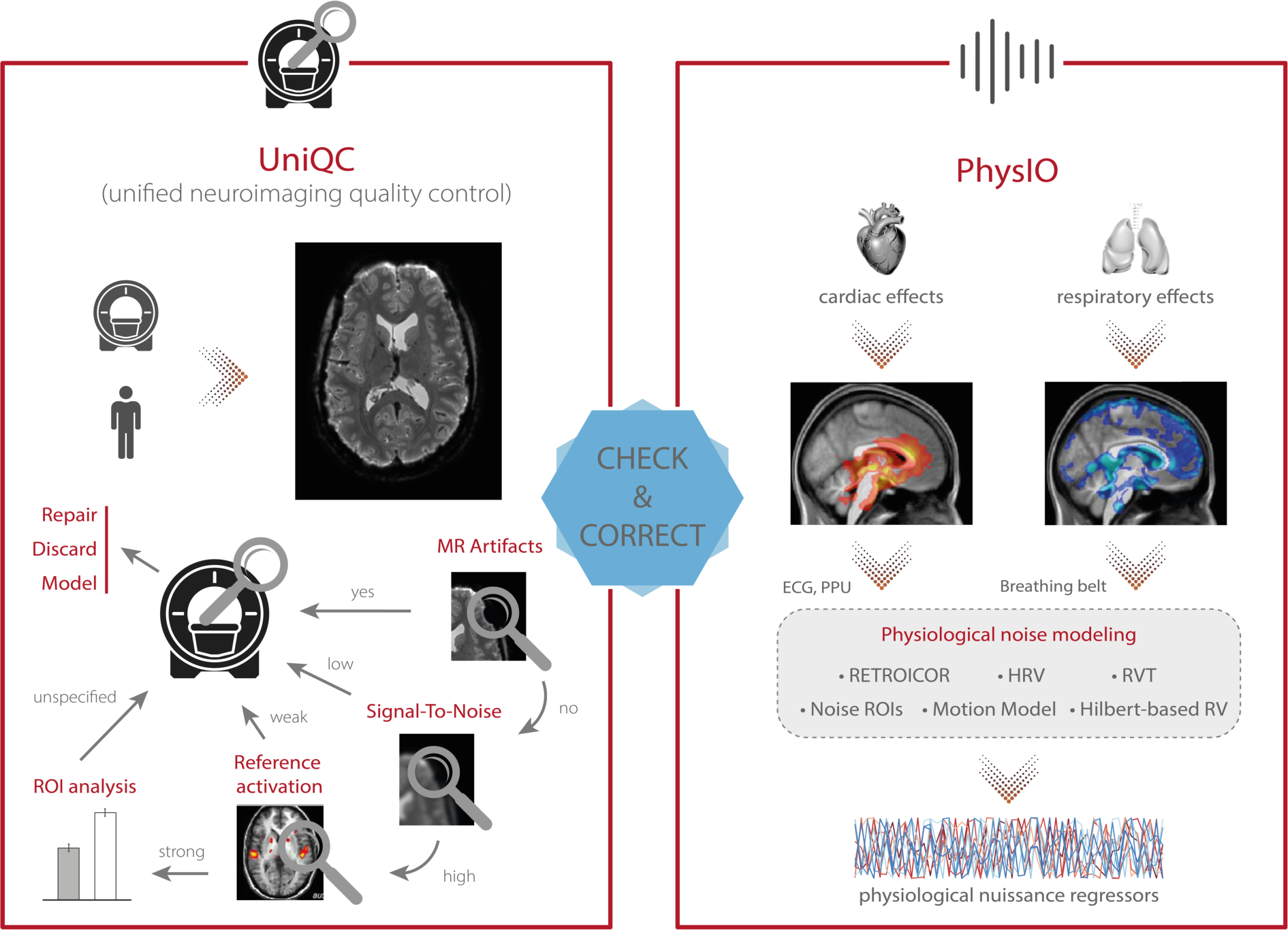
TAPAS components that aim to monitor data quality and correct for physiological confounds. (*Left*) In addition to supporting the development and optimization of MR acquisition sequences (see Figure 3), UniQC also facilitates monitoring of data quality during data acquisition in order to quickly identify potential problems in the acquisition processes which might relate to malfunctions or alterations in the scanner hardware, as well as to artifacts related to the participant (e.g., motion, physiology). (*Right*) PhysIO implements model-based physiological noise correction based on peripheral recordings of cardiac (e.g., electrocardiogram (ECG), photoplethysmographic unit (PPU)) and respiratory (e.g., breathing belt) cycle. The toolbox uses RETROICOR as well as modeling of the impact of heart rate variability (HRV) and respiratory volume per time (RVT) on the BOLD signal (e.g., using Hilbert-based respiratory volume) to derive physiological nuisance regressors that can be utilized in subsequent statistical analyses to account for physiological confounds in fMRI signals.

In principle, this fast and precise identification can lead to quality improvements in three ways (Figure 4, *left*). First, the information can be used to isolate and repair hardware malfunction (e.g., of certain coil elements) to swiftly restore quality levels for the next scan or participant. Second, the quality of the affected dataset can be increased by modeling the impact of distinct noise sources, as identified by the QC decision tree (e.g., electrostatic spike artifacts in the images or interactions between subject motion and magnetic field; Dymerska et al., 2018; Jezzard and Balaban, 1995; Jezzard and Clare, 1999). Third, selectively discarding the low-quality data only, as isolated by the customized QC interrogation, salvages quality levels for the remainder of the data (Figure 4, *left*).

Furthermore, unlike generic QC pipelines, the customization afforded by UniQC facilitates testing whether any given dataset shows a functional response relevant for the research question. For example, due to the seamless integration with other MATLAB toolboxes such as SPM, UniQC can analyze statistical maps from study-specific GLMs, as well as provide region-of-interest (ROI) statistics. With this functionality, task-fMRI performance can be validated using robust expected activation patterns as a sanity check, for example, sensory-motor activation during a learning task, before proceeding to more complex analyses. Thus, UniQC offers flexible, interactive and study-specific quality control of the image acquisition system and the imaged participant. Thanks to its modularity, UniQC can be further integrated to monitor quality throughout the preprocessing stage (section 4), independent of the concrete pipeline.

### 3.2 Physiological noise modeling

One of the main confounds in fMRI is physiological noise as it perturbs blood oxygen level dependent (BOLD) signals (Hutton et al., 2011; Kruger and Glover, 2001) – which can substantially hamper both classical fMRI analysis as well as computational modeling of the data. The two primary sources of physiological noise are the cardiac and respiratory cycles (Murphy et al., 2013). The respiratory cycle introduces confounds by distortions of the magnetic field due to the movement of the participant’s chest (Brosch et al., 2002) as well as bulk susceptibility variation in the lungs (Raj et al., 2001). Additionally, the respiratory cycle alters the pressure of blood CO2 (which is a vasodilator) over longer time periods, thereby inducing slow signal fluctuations (Birn et al., 2006). The cardiac cycle, on the other hand, modulates blood volume and vessel diameter during systole and diastole, leading to small deformations of brain tissue and brainstem displacement, causing periodic motion of the cerebrospinal fluid (Dagli et al., 1999). Furthermore, variability in heart rate induces alterations in the oxygen level in the blood (Huettel et al., 2009), and consequent low frequency signal fluctuations (Chang et al., 2009). Finally, interactions between the cardiac and respiratory cycles, such as in respiratory sinus arrhythmia (i.e., accelerated heartbeat during inhalation), induce additional non-trivial physiological fluctuations (Hirsch and Bishop, 1981).

Various physiological noise correction methods for fMRI exist, either based solely on the fMRI time series and prior assumptions of spatiotemporal noise properties, or modeling the noise from independent physiological recordings (e.g., using electrocardiogram (ECG), photoplethysmographic unit (PPU), and breathing belts) (Kasper et al., 2017; Murphy et al., 2013). Arguably, for TN/CP applications with clinical populations and pharmacological interventions, methods based on independent recordings might be preferable: In clinical populations, physiological processes that impact on the BOLD signal may differ from priors that were defined based on the general population.

Several freely available implementations for model-based physiological noise correction are available, including AFNI 3DRETROICOR (Glover et al., 2000), FSL Physiological Noise Modeling (Brooks et al., 2008), and PhLEM (Verstynen and Deshpande, 2011); however, relatively few studies have capitalized on these tools, in particular for task-based fMRI. One important practical challenge – that is exacerbated in clinical populations with less compliant subjects – is the variable data quality of the peripheral recordings which these models are based on. Reduced data quality may be due to subject motion or (partial) detachment/saturation of the peripheral devices. In most implementations, preprocessing of these recordings is minimal, and identification of the physiological cycles typically relies on peak detection provided by the MR scanner vendor (Kasper et al., 2017). Alternatively, manual intervention to correct erroneous detection is offered, hampering the development of automatized pipelines and the translation of physiological noise modeling into routine application.

The *PhysIO* toolbox (Kasper et al., 2017) in TAPAS offers methods for model-based physiological noise correction based on peripheral recordings of the cardiac (e.g., ECG, PPU) and respiratory cycles (e.g., breathing belt). PhysIO utilizes these peripheral measures to model the periodic effects of pulsatile motion and field fluctuations using RETROICOR (Glover et al., 2000). Furthermore, the toolbox accounts for end-tidal CO2 changes and heart rate-dependent blood oxygenation by convolving respiratory volume per time (RVT) and heart rate variability (HRV) with a respiratory and cardiac response function, respectively (Birn et al., 2006; Birn et al., 2008; Chang and Glover, 2009). For these methods, emphasis is placed on robust preprocessing of the input time series via reliable peak detection in low-SNR regimes, as well as a novel method for RVT estimation using the Hilbert transform (Harrison et al., 2020b). As a more data-driven alternative for noise correction, PhysIO also allows the extraction of signals from pre-defined regions of interest (“noise ROIs”) as additional confound regressors – for instance, signal related to white matter or cerebrospinal fluid (CSF). Finally, the toolbox also incorporates various strategies for correcting motion-related artifacts by implementing, for instance, the Volterra expansion confound set (Friston et al., 1996) or censoring strategies based on the framewise displacement (Power et al., 2012). A schematic illustration of the modeling process in PhysIO is provided in Figure 4, *right*. PhysIO provides both command-line operation for de-noising multiple subjects conveniently, as well as a user-friendly graphical interface within the SPM Batch Editor. Thereby, physiological noise correction can be integrated with complete fMRI preprocessing pipelines, minimizing the need for manual interventions or custom programming (Kasper et al., 2017). Additionally, PhysIO ensures robust preprocessing even for low-quality data and provides simple diagnostic tools to assess the correction efficacy in individual subjects. This renders PhysIO an accessible noise correction tool for preprocessing pipelines both in basic neuroscience studies as well as for clinical purposes.

## 4 PREPROCESSING

Data preprocessing is closely intertwined with quality control and artifact correction. Thorough preprocessing is particularly important for complex data acquired using neuroimaging techniques such as fMRI or EEG/MEG (Ashburner, 2016). To this purpose, researchers typically create ad hoc preprocessing workflows for each study individually (Carp, 2012), building upon a large inventory of available tools. Broadly speaking, preprocessing steps can be separated into two main categories: (i) preprocessed time series are derived from the original data after application of retrospective signal corrections, spatiotemporal filtering, and resampling in a target space (e.g., MNI standard space), and (ii) confound-related information in the data can be modeled or taken into account through nuisance regressors (i.e., regressors of no interest) in subsequent statistical analyses using the general linear model (GLM). These confounds may include motion parameters, framewise displacement, physiological (cardiac or respiratory) signals, or global signals (Murphy et al., 2013; Power et al., 2014).

These and additional procedures for preprocessing neuroimaging data are available in various software packages including, SPM (Friston et al., 2006), FSL (Jenkinson et al., 2012), FreeSurfer (Fischl, 2012), AFNI (Cox, 1996), or Nilearn (Abraham et al., 2014). The plethora of different preprocessing tools and workflows manifests in the absence of a current gold standard for preprocessing neuroimaging data, despite several attempts to establish best-practice guidelines (Ashburner, 2016; Gorgolewski et al., 2016; Power et al., 2017; Strother, 2006).

In an attempt towards common preprocessing standards, large-scale consortia like the HCP or the UK Biobank provide access not only to the raw data, but also to already preprocessed versions of the data. For instance, in the HCP database, researchers have access to the version of the data which have been subjected to a common minimal preprocessing pipeline (Glasser et al., 2013). However, these workflows are usually tailored towards the particular dataset’s idiosyncrasies and do not readily translate to other datasets. A first attempt towards such a universally applicable preprocessing workflow is fMRIPrep (Esteban et al., 2019), which represents a pipeline that combines tools from several of the above-mentioned software packages. fMRIPrep autonomously adapts the workflow to the present data, rendering the approach robust to data idiosyncrasies and potentially applicable to any dataset without manual intervention.

While no toolbox dedicated to data preprocessing is currently available in TAPAS, our tools from the previous step (i.e., “Conduct, Check and Correct”) integrate well with most of the third-party software packages highlighted above. For instance, PhysIO integrates seamlessly with the batch editor system of SPM to facilitate the derivation of nuisance regressors related to physiological confounds that can be utilized in GLM analyses. Similarly, UniQC is designed to integrate with SPM and other MATLAB toolboxes. Importantly, PhysIO and UniQC are also designed to integrate well with other neuroimaging software packages. For instance, PhysIO stores all physiological noise regressors in a dedicated text file which can be inputted into the first-level analyses in any software package (e.g., FSL, AFNI).

## 5 INFERENCE

Once neuroimaging and/or behavioral data have been preprocessed and artifacts have been corrected, the question arises how best to interrogate the data in order to gain insights into the functioning of the human brain and alterations thereof in disease. Concerning clinically oriented studies, it has been pointed out (Kapur et al., 2012; Stephan et al., 2015) that focusing on differences in descriptive measures – such as BOLD activation, functional connectivity patterns or task performance – between patients and healthy controls is unlikely to result in improvements of clinical practice. This is because these analyses do not provide an understanding of the symptom-producing mechanisms and they do not easily inform the development of biologically grounded clinical tests. Consequently, these measures have not yet led to routine applications in clinical practice (Filiou and Turck, 2011; Kapur et al., 2012).

To address this shortcoming, mathematical models of neuroimaging and behavioral data that capture putative physiological and cognitive disease mechanisms may represent a promising avenue. This line of thinking is at the core of clinically-oriented modeling disciplines like Computational Psychiatry (Friston et al., 2014; Huys et al., 2016; Maia and Frank, 2011; Montague et al., 2012; Stephan and Mathys, 2014; Wang and Krystal, 2014), Computational Neurology (Jirsa et al., 2016; Maia and Frank, 2011), and Computational Psychosomatics (Petzschner et al., 2017). For example, in Computational Psychiatry, a major goal is to move from the current syndromatic nosology to disease classifications based on computational assays that may improve differential diagnosis and treatment prediction for individual patients (Friston et al., 2017; Stephan et al., 2016a).

TAPAS contributes to this endeavor by providing a collection of computational tools that can be applied to neuroimaging (fMRI) or behavioral data (decisions, eye movements). All of these approaches are so-called generative models (Bishop, 2006). Generative models specify the joint probability *p*(*y, θ*|*m*) over measured data *y* and model parameters *θ*, which – according to probability theory – can be written as the product of the likelihood function *p*(*y*|*θ*, *m*), representing the probability of the data given a set of model parameters, and the prior distribution *p*(*θ*|*m*), encoding the *a priori* plausible range of parameter values. Together, likelihood and prior yield a probabilistic forward mapping from latent (hidden) states of a system (e.g., neuronal dynamics) to observable measurements (e.g., BOLD signal). Model inversion enables inference on the parameters and latent states of the system from measured data, and can be accomplished using a variety of approximate Bayesian techniques (e.g., variational Bayes, Markov chain Monte Carlo or Gaussian process optimization). Their ability to reveal latent mechanisms underneath the visible data and their natural connection to hypothesis testing through model selection procedures (Penny et al., 2004) have established generative modeling as a cornerstone of TN/CP (Frässle et al., 2018b; Stephan et al., 2017).

In brief, the generative models included in TAPAS comprise: (i) models of effective (directed) connectivity among neuronal populations, (ii) models of perception in the light of an uncertain and volatile environment, as well as (iii) models of inhibitory control. In what follows, we briefly describe the different generative models in TAPAS and highlight how each of them might be useful for clinical (neuro)modeling.

### 5.1 Generative models of neuroimaging data

Models of effective connectivity describe the mechanisms by which neuronal populations interact and how these mechanisms give rise to measured data (e.g., fMRI or EEG/MEG). By inverting these generative models, it is possible, in principle, to infer on the directed (synaptic) influences neuronal population exert on one another (Friston et al., 2013; Friston, 2011). This differs from measures of functional connectivity (e.g., Pearson’s correlation) which are essentially descriptive and undirected statistical indices. Models of effective connectivity hold particular promise for TN/CP, since global dysconnectivity has been proposed as a hallmark of various mental disorders (Deco and Kringelbach, 2014; Menon, 2011), including schizophrenia (Bullmore et al., 1997; Friston et al., 2016a; Friston and Frith, 1995; Stephan et al., 2006; Stephan et al., 2009), autism (Courchesne et al., 2007; Kennedy et al., 2006; Muller, 2007), and depression (Greicius et al., 2007; Mayberg, 1997; Wang et al., 2012).

A frequently used generative modeling framework for inferring effective connectivity from neuroimaging data is dynamic causal modeling (DCM; Friston et al., 2003). In brief, DCM describes changes in neuronal activity as a function of the directed interactions among neuronal populations and experimental manipulations that can perturb the system. DCM was initially introduced for fMRI (Friston et al., 2003) and later extended to electrophysiological data (David et al., 2006). Comprehensive reviews on DCM can be found elsewhere (e.g., Daunizeau et al., 2011; Friston et al., 2013; Moran et al., 2013; Stephan et al., 2010). DCM is freely available as part of SPM and has found widespread application. For example, with regard to clinical applications, DCM has been used to study schizophrenia (Brodersen et al., 2014; Deserno et al., 2012; Dima et al., 2009; Lefebvre et al., 2016; Li et al., 2017), autism (Grèzes et al., 2009; Radulescu et al., 2013), and depression (Almeida et al., 2009; Schlösser et al., 2008). Despite these promises, the classical DCM approach is also subject to several limitations – which may become particularly relevant in the context of TN/CP, where the goal is to develop computational assays that inform prediction of clinical trajectories and treatment responses in individual patients. In what follows, we highlight some of these limitations and outline how the methodological advances in DCM included in TAPAS aim to address these challenges.

#### 5.1.1 Global optimization

When translating computational advances like DCM into computational assays, the robustness of the inference procedure and the reliability of the parameter estimates become paramount (Woolrich and Stephan, 2013). Standard model inversion in DCM rests on variational Bayes under the Laplace approximation (VBL; Friston et al., 2007) which is computationally efficient, yet subject to several limitations (Daunizeau et al., 2011): First, VBL rests on maximizing the negative free energy (which serves as a lower bound approximation to the log model evidence) using gradient ascent and is thus inherently susceptible to local maxima if the objective function is multimodal. Second, even when the global maximum is found, the distributional assumptions (i.e., Laplace and mean-field approximations) might not be justified, potentially rendering the approximate posterior distribution a poor representation of the true posterior. Third, when the distributional assumptions of the Laplace approximation are violated, the negative free energy is no longer guaranteed to represent a lower bound on the log model evidence (Wipf and Nagarajan, 2009).

Sampling-based model inversion schemes, typically based on Markov chain Monte Carlo (MCMC) methods, do not require any distributional assumptions about the posterior and are guaranteed to be asymptotically exact (i.e., converge to the global extremum in the limit of infinite samples). This renders sampling-based methods an appealing alternative to VBL. However, they come at the cost of other challenges: First, sampling-based routines are computationally expensive. Second, convergence is only guaranteed in the limit of infinite samples; detecting convergence in practice thus rests on heuristics. Third, unlike VBL, sampling-based methods do not readily provide an estimate of the (log) model evidence, but require additional strategies, which further aggravate the computational burden. For instance, the current gold standard for sampling-based estimates of the model evidence, thermodynamic integration (TI; Calderhead and Girolami, 2009; Kirkwood, 1935; Lartillot and Philippe, 2006), requires running multiple MCMC chains at different “temperatures” (i.e., at different positions along a path from prior to posterior). Until recently, these reasons have been prohibitive for the use of sampling-based model inversion for DCMs.

The *massively parallel dynamic causal modeling* (mpdcm) toolbox (Aponte et al., 2016) implemented in TAPAS renders sampling-based model inversion in the context of DCM for fMRI computationally feasible (Figure 5, *top*). This is achieved by exploiting the power of graphics processing units (GPUs) for the evaluation of the likelihood function, which represents the computationally most expensive operation, as it requires integration of differential equations in the neuronal and hemodynamic models. Importantly, mpdcm even makes the evaluation of the model evidence via thermodynamic integration computationally feasible. In a recent preprint, Aponte et al. (2020b) demonstrated that TI provides more accurate and robust estimates of the model evidence than VBL, while computational demands are kept at a moderate level.

**Figure 5.**
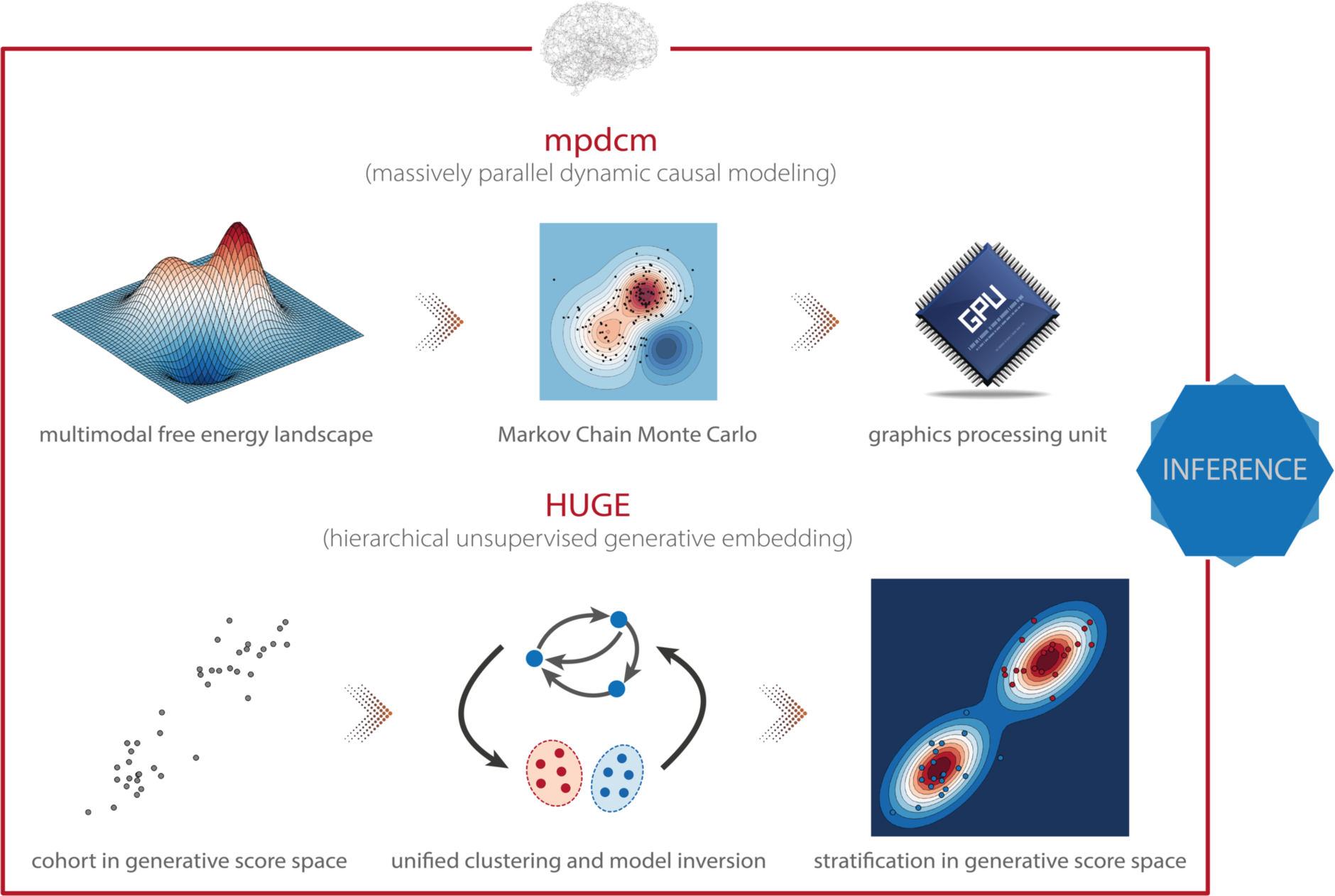
TAPAS components that implement generative models of neuroimaging data (I). (*Top*) Massively parallel dynamic causal modeling (mpdcm) renders sampling-based model inversion computationally feasible by exploiting graphics processing units (GPUs). This allows one to obtain more faithful results in the presence of a multimodal optimization landscape. (*Bottom*) Hierarchical unsupervised generative embedding (HUGE) combines the inversion of single-subject DCMs and the clustering of participants into mechanistically homogenous subgroups within a single generative model.

Beyond the mpdcm toolbox, which is designed to support DCM for fMRI, other gradient-free and gradient-based MCMC sampling schemes have also been introduced to DCM for electrophysiological data (Sengupta et al., 2015, 2016). However, these tools have not yet been made publicly available.

#### 5.1.2 Empirical Bayes for DCM

Another challenge concerns the specification of prior distributions in DCM, which have been found to profoundly impact the posterior estimates and their reliability (Frässle et al., 2015). Notably, in the context of hierarchical Bayesian models, there is a principled way of estimating priors by exploiting measurements from multiple subjects: empirical Bayes (EB; Banerjee et al., 2015; Efron and Morris, 1973; Kass and Steffey, 1989). In brief, in EB, the posterior density at any given level is constrained by the level above. For instance, in a two-level hierarchical model, observed data *y* = {*y*_!_, *y*_”_, …, *y*_#_} are assumed to be generated from a set of latent (hidden) parameters *θ* = {*θ*_!_, *θ*_”_, …, *θ*_#_} according to the likelihood *p*(*y*|*θ*, *m*). In turn, the parameters *θ* are considered to represent samples from a population density *p*(*θ*|*η*, *m*), where *η* refers to the hyperparameters (Bishop, 2006; Gelman et al., 2004). Consequently, under the hierarchical structure of a multi-subject or mixed-effects model, inference on the single-subject level is constrained by the group-level information. These constraints are then referred to as empirical priors since they are informed by the empirical data (of the entire group). A special case of EB is referred to as parametric empirical Bayes (PEB), where the hyperparameters *η* are approximated using the maximum likelihood estimate or a moment expansion, which allows one to express the hyperparameters in terms of the empirical mean and variance (Maritz and Lwin, 1989). One particular variant of PEB is the Gaussian-Gaussian model, where single-subject data are assumed to be generated by adding Gaussian noise to the group mean (Friston et al., 2016b).

The *hierarchical unsupervised generative embedding* (HUGE) toolbox contained in TAPAS implements EB in the context of DCM for fMRI (Raman et al., 2016; Yao et al., 2018). HUGE combines the inversion of single-subject DCMs and the clustering of subjects into mechanistically homogenous subgroups into a single generative model (Figure 5, *bottom*). This is achieved by combining the nonlinear DCMs at the individual level with a mixture-of-Gaussians clustering model at the hierarchically higher level (Bishop, 2006; Gelman et al., 2004). In other words, HUGE assumes that each individual from a population of *N* subjects belongs to one of at most *K* subgroups or clusters. The DCM parameters *θ* for all subjects from one cluster *k* are then assumed to be normally distributed with distinct mean μ_*k*_ and covariance matrix Σ_*k*_. This cluster-specific normal distribution effectively means that different prior distributions apply over subjects, depending on which subgroup they belong to, and that these priors are learned from the data (i.e., subgroup-specific EB). Hence, in principle, the framework is capable of stratifying heterogenous spectrum disorders, as defined by DSM/ICD, into subgroups that share common pathophysiological mechanisms (for more details, see below). Importantly, HUGE also implements “pure” EB by fixing the number of clusters to one and merely exploiting the hierarchical dependencies in the data. This effectively switches off the clustering model. The utility of this mode of operation has been demonstrated in simulations by Yao et al. (2018), highlighting the expected shrinkage effect (reduced variability) of the posterior parameter estimates towards the population mean observed in EB (Bishop, 2006). Parameter estimation in HUGE can be performed by employing either a sampling-based MCMC inversion scheme (Raman et al., 2016; Yao and Stephan, 2020), which is asymptotically exact yet (relatively) slow, or a VB implementation (Yao et al., 2018), which is computationally more efficient yet might be vulnerable to local extrema. Notably, at the moment, only the VB implementation of HUGE is available in TAPAS. The sampling-based variant will be published as part of an upcoming release of the toolbox.

#### 5.1.3 Whole-brain effective connectivity analysis

Apart from the computational and statistical challenges mentioned above, a conceptual concern is that DCMs are typically restricted to relatively small networks in order to keep model inversion computationally feasible. While this may be advantageous in some cases by enforcing a theory-driven analysis of high-dimensional and noisy fMRI data, it can also represent a limiting factor. Specifically, many cognitive processes, as well as the “resting state” (i.e., unconstrained cognition in the absence of experimental manipulations), engage a widespread network that cannot be captured faithfully by a handful of nodes. Furthermore, in the context of Computational Psychiatry, putative pathophysiological processes underlying various mental disorders have been linked to global (large-scale) alterations of functional integration in brain networks (e.g., Courchesne et al., 2007; Friston et al., 2016a; Friston and Frith, 1995; Greicius et al., 2007; Kennedy et al., 2006; Mayberg, 1997; Stephan et al., 2006; Stephan et al., 2009). This calls for the development of computational models that are capable of inferring effective (directed) connectivity in whole-brain networks (Menon, 2011).

*Regression dynamic causal modeling* (rDCM; Frässle et al., 2018a, 2017) represents a recent variant of DCM that renders model inversion extremely efficient. This is achieved by converting the numerically costly estimation of coupling parameters in differential equations of a linear DCM in the time domain into a Bayesian linear regression model in the frequency domain (Figure 6, *top*). Under a suitably chosen mean-field approximation, analytically solvable VB update equations can be derived for this model. The ensuing computational efficiency allows rDCM to scale gracefully to large-scale networks that comprise hundreds of regions. Furthermore, rDCM has recently been augmented with sparsity constraints to automatically prune fully connected networks to an optimal (in terms of maximal model evidence) degree of sparsity (Frässle et al., 2018a). This is achieved by introducing binary indicator variables into the likelihood function, which essentially serve as feature selectors. For this generative model, comprehensive simulation studies demonstrated the face validity of rDCM with regard to model parameter and model architecture recovery. Furthermore, we have provided initial demonstrations of the construct validity of the approach in applications to empirical data. For instance, using ultra-high field (7T) fMRI data from a simple hand movement paradigm with the known relevant connections, we demonstrated that rDCM inferred plausible effective connectivity patterns in whole-brain networks with more than 200 regions (Frässle et al., 2021). Furthermore, we have recently demonstrated that rDCM can not only be applied to task-based, but also to resting-state fMRI data (Frässle et al., 2020a). Notably, inversion of whole-brain models with rDCM is computationally highly efficient on standard hardware: even for whole-brain networks with more than 200 regions, it takes only a couple of minutes for fixed network architectures, and a few hours when pruning fully connected networks.

**Figure 6.**
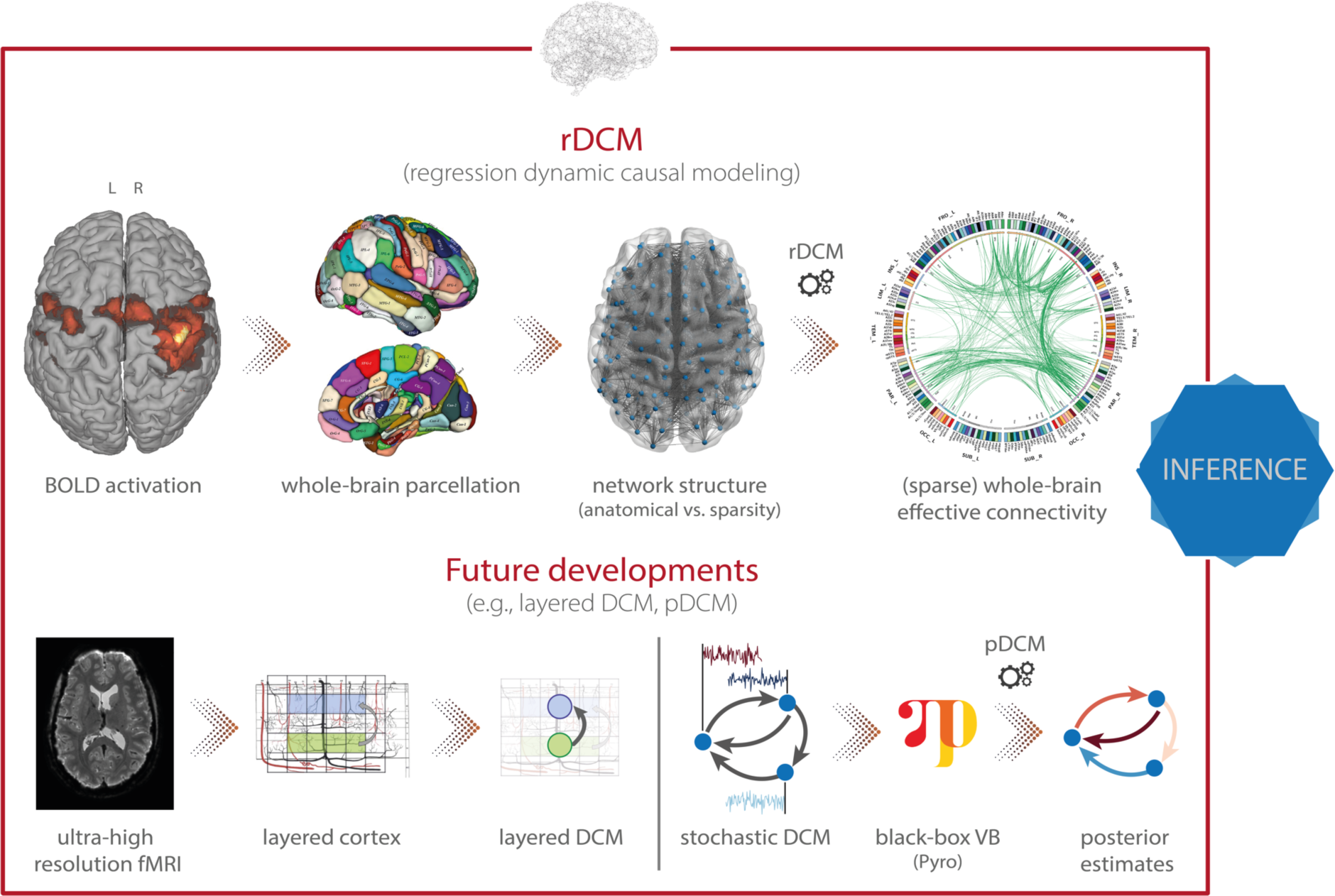
TAPAS components that implement generative models of neuroimaging data (II). (*Top*) Regression dynamic causal modeling (rDCM) is a novel variant of DCM for fMRI that scales gracefully with the number of nodes and thus makes whole-brain effective connectivity analyses feasible. (*Bottom*) Layered DCM (l-DCM) and pDCM as future developments of DCM, representing tools that will be included in TAPAS in one of the next upcoming releases. Parts of figure reproduced with permission from (Heinzle et al., 2016b), Copyright 2016 Elsevier, and (Frässle et al., 2021), Copyright 2020 Elsevier.

#### 5.1.4 Future developments

Besides the toolboxes mentioned above, which are already part of TAPAS, additional variants of DCM for fMRI will be released soon. Specifically, this includes: (i) layered dynamic causal modeling (layered DCM; Heinzle et al., 2016b), and (ii) pDCM, a Python-based DCM implementation focused on amortized inference of stochastic DCMs using novel probabilistic-programming techniques (Harrison et al., *in preparation*). Here, we briefly outline these two advances.

First, layered DCM addresses challenges in effective connectivity analyses that become relevant when moving towards high-resolution fMRI measurements at the sub-millimeter scale (Figure 6, *bottom left*). This allows differentiating BOLD signals from different cortical layers, an important aspect for testing desiderata of modern theories of brain structure and function. Specifically, prominent “Bayesian brain” theories like predictive coding (Friston, 2005a; Rao and Ballard, 1999) postulate that supragranular and infragranular cortical layers convey different signals via their efferent cortico-cortical connections. Testing these theories might not only further our understanding of the functioning of the human brain in health, but also has important implications for delineating pathophysiological processes in disease.

However, for layered fMRI data, the spatial layout of cortical blood supply – in particular, the venous blood draining back from lower layers to the cortical surface – confounds responses in different layers and thus renders interpretations non-trivial. Accounting for such draining effects is thus important, yet not readily possible within contemporary hemodynamic models, such as the Balloon model currently implemented in DCM (Buxton et al., 1998; Friston et al., 2000; Stephan et al., 2007) or more recent hemodynamic models that strive for increased biological plausibility (Havlicek et al., 2015). Layered DCM addresses this limitation by extending the classical Balloon model with a phenomenological description of blood draining effects. This rests on including a delayed coupling of the relative blood volume and deoxyhemoglobin concentration across layers. For this framework, Heinzle et al. (2016b) demonstrated the face validity using simulation studies, as well as the practical utility in an application to empirical fMRI data from a simple visual paradigm. While a detailed dynamic model of layered blood flow effects was published recently (Havlicek and Uludag, 2020), this model requires far more parameters and does render inversion on empirical data more difficult.

Second, a novel inversion scheme for stochastic DCMs will be included in TAPAS, leveraging recent breakthroughs in black-box variational inference (Harrison et al., *in preparation*). In brief, the approach makes use of optimization algorithms from deep learning software packages to allow inference of general probabilistic models (Figure 6, *bottom right*). For inference, one directly infers the neuronal and hemodynamic parameters, but temporal convolutional neuronal networks are used to amortize the inference of the neuronal states themselves. This allows inferring the hidden (neuronal) states for any length of input data while keeping the number of parameters fixed (instead of growing linearly with the number of time points). The advantage of using black-box variational inference algorithms is that the framework is highly flexible, allowing for easy modifications and extensions of the underlying generative model without any changes to the inference machinery. This renders the model very promising for clinical applications where generative models might need to be tailored towards specific diseases.

### 5.2 Generative models of behavioral data

Neuroimaging provides functional readouts from disease-relevant neural circuits and thus delivers data for models of pathophysiology. However, these data and models are usually not suitable for drawing direct conclusions about cognition and its disturbances. By contrast, behavioral data can be used for inference on an agent’s internal processes at the algorithmic (information processing) level. Importantly, acquisition of behavioral data is often easier, cheaper and more patient-friendly than neuroimaging data, and computational models of behavior thus hold great promise for establishing clinically useful computational assays – on their own or in combination with neuroimaging data (Browning et al., 2020; Stephan and Mathys, 2014). At present, TAPAS contains two different generative models of behavioral data which will be discussed next: (i) the Hierarchical Gaussian Filter (HGF; Mathys et al., 2011), and (ii) the Stochastic Early Reaction, Inhibition and late Action model (SERIA; Aponte et al., 2017).

#### 5.2.1 Hierarchical Gaussian Filter (HGF)

The *Hierarchical Gaussian Filter* (HGF; Mathys et al., 2011) is a hierarchical Bayesian framework for individual learning under the various kinds of uncertainty which arise in realistic nonlinear dynamic systems (e.g., perceptual uncertainty, environmental volatility). Importantly, the hierarchy implemented in the HGF is not to be confused with the hierarchy that was discussed in the context of empirical Bayes and, more specifically, HUGE. While in empirical Bayes, the hierarchy (typically) refers to a multi-subject or mixed-effects structure where the levels represent single-subject and group-level information, the HGF implements a hierarchy in which the levels represent the temporal evolution of latent states. More specifically, it consists of two parts: a generative model, the HGF-GM, which describes the stochastic evolution of the nonlinearly coupled hidden states of a dynamic system; and the HGF proper, a set of deterministic update equations resulting from the variational inversion of the HGF-GM. The HGF proper contains the Kalman filter as a special case, but is also suited for filtering inputs generated by nonlinear environments. Combined with an observation model, the HGF proper represents a particular implementation of the “observing the observer” framework developed by Daunizeau et al. (2010a; 2010b). This framework is based on the separation of two model components: (i) a perceptual model which describes an agent’s inference on the environment (in this case the HGF proper), and (ii) a response model which describes how inferred latent states of the agent translate into the agent’s observed actions, such as, decisions or responses (Mathys et al., 2014).

In the HGF, the perceptual model takes the form of a hierarchical Bayesian model where the temporal evolution of states at any level (except the first) are represented as Gaussian random walks or first-order autoregressive processes (Figure 7, *top*). Importantly, the step size of each walk (i.e., the variance of the Gaussian distribution) depends on the state at the next higher level. This coupling between levels is controlled by subject-specific parameters that shape the influence of uncertainty on learning. Under a VB approximation, one can derive efficient trial-by-trial update equations for this model that describe the agent’s belief updating. Importantly, these update equations rest on precision-weighted prediction errors (PE) at different levels of the hierarchy. In other words, the HGF tracks an agent’s expression of (approximate) Bayesian learning in the presence of uncertainty under the assumption that the brain continuously updates a hierarchical generative model of sensory inputs, with PEs serving as the teaching signal. This perceptual model is then combined with a response model (e.g., unit-square sigmoid or softmax function; although a wide range of different response models is available) that links the agent’s current estimates of the latent states to observed actions, such as motor or physiological responses (Mathys et al., 2014). In combination with priors on the model parameters, this specifies a full generative model of observed responses that is inverted using maximum-a-posteriori (MAP) estimation. In summary, the HGF provides a generic approximation to subject-specific forms of hierarchical Bayesian learning^3^. For applications in computational psychiatry, parameter estimates can be used to characterize perceptual inference and decision-making in specific disorders (e.g., Lawson et al., 2017; Powers et al., 2017); alternatively, and perhaps even more frequently, the estimated trajectories of precision-weighted

**Figure 7.**
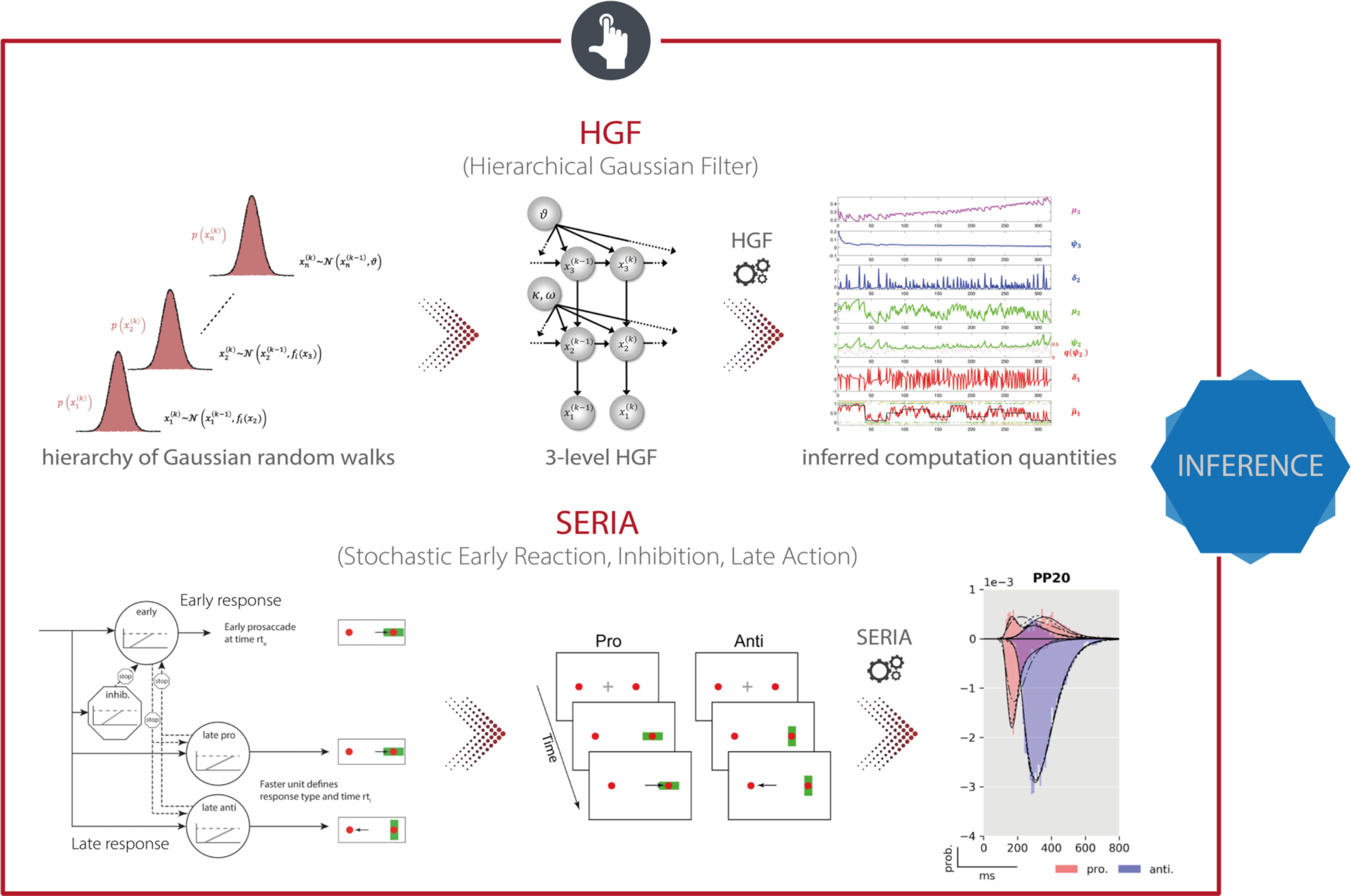
TAPAS components that implement generative models of behavioral data. (*Top*) The Hierarchical Gaussian Filter (HGF) is a hierarchical Bayesian model for individual learning under different forms of uncertainty (e.g., perceptual uncertainty, environmental volatility). (*Bottom*) The Stochastic Early Reaction, Inhibition, and late Action (SERIA) model represents a computational model of an agent’s behavior during the antisaccade task by modeling early reflexive and late intentional eye movement via two interacting race-to-threshold processes.

PEs are used in trial-by-trial analyses of fMRI and EEG data (e.g., Cole et al., 2020; Iglesias et al., 2013; Weber et al., 2020). The HGF can also be applied to other time series than behavioral ones. In an example from TN/CP, Brazil et al. (2017) applied an HGF directly to BOLD signal time series from an fMRI experiment.

#### 5.2.2 Stochastic Early Reaction, Inhibition and late Action (SERIA) model

Eye movements represent a potentially very interesting functional readout for TN/CP. In addition to the experimental ease with which many data points can be measured, eye movements are disturbed in numerous psychiatric conditions (Heinzle et al., 2016a; Hutton and Ettinger, 2006; Rommelse et al., 2008). An experimental paradigm that has been used frequently in this context is the antisaccade task (Hallett, 1978) where participants are asked to suppress a reactive eye movement towards a visual cue and concurrently perform a saccade in the opposite direction (antisaccade). This task is of relevance for clinical applications since it has been widely used to study psychiatric and neurological diseases (Hutton and Ettinger, 2006). Most prominently, the antisaccade task is hampered in schizophrenia (Curtis et al., 2001; Fukushima et al., 1988) where an elevated error rate has been proposed as an intermediate phenotype or endophenotype of the disease (Reilly et al., 2014).

TAPAS comprises a computational model of an agent’s behavior during the antisaccade task – the *Stochastic Early Reaction, Inhibition and late Action* (SERIA) model (Aponte et al., 2017). Specifically, in order to model error rates and reaction times during the task, SERIA postulates two interacting processes (Figure 7, *bottom*): (i) a fast GO/NO-GO race between a prepotent response (prosaccade) towards the visual cue and a signal to cancel this erroneous action, and (ii) a slow GO/GO race between two units encoding the cue-action mapping, accounting for slow voluntary saccades. The parameters of this model, which are estimated using a sampling-based hierarchical Bayesian scheme, are sensitive to dopaminergic and cholinergic manipulations and were found to allow for out-of-sample predictions about the drug administered to an individual with 70% accuracy (Aponte et al., 2020a).

#### 5.2.3 Future developments

As for the generative models of neuroimaging data, extensions to the behavioral models will also be released as part of TAPAS in the future. In particular, an extension of the HGF is currently under development which aims to embed the classical HGF within an empirical Bayesian (EB) scheme (Mathys et al., *in preparation*). This hierarchical (in the sense of an EB scheme) version of the HGF combines HGFs at the individual level with a layer that represents group effects. Similar to HUGE, this formulation affords a principled way of estimating (empirical) priors from the data, effectively constraining single-subject estimates by group-level information. Inference in the H2GF rests on sampling-based MCMC methods that provide not only an estimate of individual model parameters, but also an approximation to the model evidence via thermodynamic integration (or other suitable techniques).

## 6 CLINICAL APPLICATION

Computational assays are developed with the goal to solve concrete clinical problems. Here, we briefly consider 3 types of problems: (i) differential diagnosis, (ii) stratification of spectrum disorders (i.e., identification of subgroups), and (iii) prediction of clinical trajectories or treatment responses (Stephan et al., 2015). Different computational strategies are available to address each of these challenges.

First, differential diagnosis can be formalized as hypothesis testing, which – in a Bayesian framework – is equivalent to Bayesian model selection (BMS; Bishop, 2006; Gelman et al., 2004) where different hypotheses (models) are compared in the light of observed neuroimaging and/or behavioral data. Specifically, by formalizing competing pathophysiological and/or psychopathological theories in terms of distinct models, differential diagnosis boils down to assessing the relative plausibility of these models (Stephan et al., 2017). This rests on comparing the model evidence which represents a principled metric of model goodness that trades off accuracy and complexity. Importantly, all generative models included in TAPAS provide an approximation to the model evidence and thus lend themselves, in principle, to the idea of hypothesis testing as a formal way to compare the plausibility of alternative disease mechanisms.

Second, prediction of clinical trajectories and treatment outcome as well as stratification of spectrum disorders can be achieved by means of supervised and unsupervised generative embedding (GE; Shawe-Taylor and Cristianini, 2004), respectively. In brief, the key idea of GE is to perform (un)supervised learning in a feature space that is spanned by the posterior estimates obtained from a generative model fitted to the data. In other words, the generative model serves as a theory-driven dimensionality reduction device which projects the high-dimensional and noisy data onto neurobiologically meaningful parameters that span a low-dimensional and interpretable space for (un)supervised learning. GE frequently yields more accurate results than conventional ML (Brodersen et al., 2014; Brodersen et al., 2011; Frässle et al., 2020b), likely because the generative model separates signal (reflecting the process of interest) from (measurement) noise more efficiently than other methods. In what follows, we outline toolboxes included in TAPAS that can be used for supervised and unsupervised GE.

### 6.1 Classification and prediction

Typically, GE is implemented in terms of a two-step procedure: (i) generative models of measured data are inverted for each subject individually, and (ii) the summary statistics of the posterior estimates (e.g., the maximum-a-posteriori estimates) are used for supervised (classification, regression) or unsupervised (e.g., clustering) learning.

Here, we first focus on supervised GE as a formal way of performing differential diagnosis (via classification) or outcome prediction. To this end, TAPAS comprises the *Generative Embedding* (GE) toolbox, Python-based software that facilitates the generative embedding framework and allows for exploration and visualization of classification performance. The toolbox is a wrapper around scikit-learn (Pedregosa et al., 2011), with the goal of providing a set of convenient functions and sensible defaults that form a suitable starting point for generative embedding analyses. The GE toolbox can use posterior parameter estimates from any of the generative models mentioned above as input features, and perform binary or multi-class classification. The toolbox utilizes logistic regression as its default classifier because it represents a simple linear model that protects against overfitting, is (relatively) simple to interpret, and the ensuing class probabilities are useful for interpreting classifier outputs. Furthermore, the toolbox implements a repeated k-fold cross-validation as the default procedure for both model selection (i.e., hyperparameter tuning) and model validation (i.e., estimating out-of-sample performance), including allowing for within-fold confound correction (Snoek et al., 2019). This choice is motivated by the fact that, in comparison to leave-one-out cross-validation, k-fold cross-validation has a lower variance and is therefore less prone to overfitting (Varoquaux et al., 2017). Finally, significance testing of classification performance is done by default using permutation tests as they provide an unbiased estimate of error variance. This is in contrast to parametric tests (e.g., binomial confidence intervals, McNemar’s test) which typically underestimate variance and are therefore overconfident (Varoquaux, 2018).

### 6.2 Stratification of heterogenous psychiatric disorders

The goal of stratifying heterogenous disorders is to identify subgroups that share common pathophysiological or psychopathological mechanisms. The increased homogeneity in terms of underlying disease mechanisms increases the power of clinical trials and enhances predictions of clinically relevant outcomes (Stephan et al., 2015). One way to achieve this goal is by using posterior parameters from a generative model for unsupervised learning (e.g., clustering); see Brodersen et al. (2014) for an example. An alternative to this two-step procedure is implemented in HUGE, a toolbox we already discussed in the context of generative models of neuroimaging data (Raman et al., 2016; Yao et al., 2018). Specifically, HUGE casts unsupervised GE as a single hierarchical generative model that simultaneously describes individual data generation and assigns participants to clusters (Figure 5, *bottom*). Unifying these two steps has a couple of conceptual advantages: (i) the hierarchical nature of the model allows learning prior distributions from the data (i.e., empirical Bayes), (ii) model inversion at the single-subject level is regularized by (cluster-specific) group results, and (iii) clustering takes the uncertainty about individual connectivity parameter estimates into account.

## 7 TAPAS IN ACTION

Finally, we briefly discuss a few selected examples from previous work that made use of different toolboxes from TAPAS. In particular, here we focus on studies that investigate pathophysiological/pathocomputational mechanisms and/or explore the role of neuromodulatory transmitter systems in these processes. The latter are a particularly prominent topic for clinical applications of generative models because the majority of available pharmacotherapeutic approaches in psychiatry targets synthesis, metabolism or receptors of neuromodulatory transmitters.

In order to non-invasively infer upon the status of neuromodulatory systems (e.g., dopamine, acetylcholine, serotonin, noradrenaline), various studies have combined experimental manipulations of different neuromodulatory systems with generative modeling of neuroimaging or behavioral data. In a first step, Iglesias et al. (2013) have provided evidence for hierarchical belief updating during a sensory associative learning task under volatility and without rewards. Hierarchical belief updating via PEs plays a central role in “Bayesian brain” theories, such as predictive coding (Friston, 2009; Rao and Ballard, 1999). Iglesias et al. (2013) utilized the HGF to infer upon subject-specific trajectories of precision-weighted PEs at different levels of the hierarchy which were then used in a GLM of fMRI data. They found that low-level PEs, encoding the mismatch between prediction and actual visual stimulus outcome, were reflected by widespread BOLD activity in visual and supramodal areas, but also in the midbrain. Conversely, high-level PEs, encoding the mismatch between prediction and actual stimulus probabilities, were reflected by BOLD activity in the basal forebrain. Midbrain and basal forebrain contain dopaminergic and cholinergic neurons, respectively, suggesting that (i) dopaminergic midbrain neurons might signal PEs unrelated to reward and (ii) cholinergic neuron activity in the basal forebrain might reflect PEs about probabilities and may thus relate to “expected uncertainty” (Yu and Dayan, 2005). Although a subsequent pharmacological study in human volunteers using the same paradigm did not support this notion (Iglesias et al., 2020), the finding that midbrain activity may reflect reward-unrelated prediction errors has since been replicated in several animal (Stalnaker et al., 2019; Takahashi et al., 2017) and human studies (Suarez et al., 2019).

Importantly, neural correlates of computational quantities can not only be detected in fMRI data, but can also be found in EEG signals, where the superior temporal resolution allows for characterizing their precise temporal dynamics. For instance, Weber et al. (2020) related HGF estimates to single-trial EEG data from participants who received ketamine in a placebo-controlled, double-blind, within-subject fashion. The authors demonstrated that PE-related activity was found in a temporal order consistent with hierarchical Bayesian theory. Additionally, they observed a significant impact of ketamine on the high-level PE about transition probabilities. Focusing on behavior only, further evidence has been provided for associations between computational quantities and the status of neuromodulatory systems. For instance, Vossel et al. (2014) perturbed the cholinergic system using pharmacological interventions, and utilized the HGF to demonstrate that this led to an increase in the rate of belief updating about cue validity during a modified Posner’s task. Similarly, Marshall et al. (2016) utilized pharmacological interventions in combination with the HGF to characterize the influence of noradrenergic, cholinergic and dopaminergic antagonists on individual estimates of uncertainty during a probabilistic serial reaction time task. The authors identified different roles for the different neuromodulatory systems, linking noradrenaline to unexpected uncertainty, acetylcholine to environmental uncertainty, and dopamine to uncertainty representations for fast, adaptive responses. Finally, Aponte et al. (2020a) demonstrated that computational quantities sensitive to neuromodulatory processes can also be derived from generative models of reflexive eye movements. Specifically, the authors conducted a double-blind placebo-controlled pharmacological study and found that computational quantities derived from an antisaccade task using the SERIA model can distinguish between dopaminergic and cholinergic effects on action selection and inhibitory control, allowing for out-of-sample predictions about the drug administered with 70% accuracy. In summary, both the HGF and SERIA comprise computational quantities that are sensitive to the functional status of different neuromodulatory systems.

Beyond questions of pathophysiology and pharmacology, tools from TAPAS have also been used to characterize clinical populations. For instance, Powers et al. (2017) studied conditioned auditory hallucinations in four groups of people who differed both in their voice-hearing and treatment-seeking statuses. Utilizing the HGF to infer upon the participants’ individual beliefs, the authors demonstrated that the weighting of prior beliefs was significantly larger in people with hallucinations than their non-hallucinating counterparts. This is consistent with the hypothesis that, in the context of a Bayesian brain, hallucinations may be explained by overly strong priors (Friston, 2005b). Focusing on patients with autism spectrum disorder (ASD), Lawson et al. (2017) utilized the HGF to provide evidence that ASD patients tend to overestimate volatility in the face of environmental changes. This leads to reduced learning about unexpected (surprising) events, which might serve as an explanation for the typical insistence on sameness and intolerance of change in ASD patients (Pellicano and Burr, 2012). Finally, Cole et al. (2020) applied HGF estimates from an associative learning task to characterize brain responses to precision-weighted PEs in individuals at clinical high risk (CHR) for psychosis. Compared to a healthy control group, CHR individuals showed enhanced PE responses in several (particularly prefrontal) regions, consistent with the prediction from the dysconnection hypothesis of schizophrenia (Friston et al., 2016a; Stephan et al., 2009) that (proneness to) psychosis is characterized by abnormal precision-weighted PE signaling in cortex. Furthermore, prefrontal PE activity was correlated with clinical status.

TAPAS tools that implement generative models of neuroimaging data have also been applied to clinical populations – although this is still rare. For instance, Yao et al. (2018) applied HUGE to an fMRI dataset comprising aphasic patients (with a lesion in the left frontal and/or temporal cortex) and healthy controls (Schofield et al., 2012) for an initial demonstration of the potential clinical utility of the model for patient stratification. In brief, the authors demonstrated that HUGE correctly identifies two clusters in the dataset, which mapped almost perfectly onto aphasic patients and healthy controls, yielding a balanced purity of 95.5%. While it is important to emphasize that diagnosing patients with aphasia does not yet represent a truly meaningful clinical problem, it demonstrates the practical utility of HUGE for stratification in a scenario where ground truth is known. Furthermore, regression DCM has been used to study alterations in whole-brain effective (directed) connectivity between psychotic patients, their first-degree relatives, as well as matched healthy controls (Frässle et al., *in preparation*). The authors demonstrate that patients showed distinctly different whole-brain connectivity patterns from healthy controls and first-degree relatives, and that the connectivity patterns allow for significant discrimination at the individual level.

Overall, the above studies illustrate the potential of generative models of behavioral and neuroimaging data for clinical applications. However, these studies do not yet implement the kind of end-to-end analysis pipeline that we outlined, at the beginning of the article, as a basis for future computational assays. Instead, the above studies simply used selected components from TAPAS at a time. Having said this, recent work by Harrison and colleagues comes close to the kind of end-to-end pipeline highlighted above, combining multiple components from TAPAS (Harrison et al., *in preparation*). In brief, the authors aimed to investigate interoception and how anxiety relates to the perception of internal bodily states. To this end, Harrison and colleagues employed two paradigms available in TAPAS Tasks, namely the Filter Detection (FD) and Breathing Learning (BL) task. The FD task revealed differences in sensitivity to breathing perception and altered interoceptive metacognitive bias between low-anxiety and moderate-anxiety healthy controls. Furthermore, for the BL task, the authors acquired fMRI data using a high-field 7T MR scanner. PhysIO was employed for physiological noise correction based on measurements of cardiac and respiratory cycles. fMRI then underwent thorough preprocessing and artefact removal by combining tools from various software packages, including FSL and SPM. Brain activity coupled with dynamic changes in bodily states was then modelled using subject-specific trajectories of predictions and PE which have been inferred utilizing the HGF toolbox. This revealed the anterior insula to be associated with both interoceptive predictions and PEs, where the former was also differentially expressed in the low and moderate anxiety groups.

While this moves towards the kind of end-to-end analysis pipeline that we outline, it is important to note that the two groups tested by Harrison and colleagues do not represent clinical groups (but were recruited from the healthy population) and the study thus lacks the final module of the aforementioned pipeline (i.e., Clinical Application; see Figure 2). Furthermore, it is important to keep in mind that none of the studies mentioned above is yet of any direct clinical utility, in the sense that they do not address a practical clinical question, such as differential diagnosis or predicting outcomes/clinical trajectories. The latter in particular requires data from prospective studies that include information about future clinical outcomes – a critical condition for validating computational assays (Stephan and Mathys, 2014). Unfortunately, so far, these datasets are rare.

## 8 CONCLUSION

In this article, we have described the **T**ranslational **A**lgorithms for **P**sychiatry-**A**dvancing **S**cience (TAPAS) software package, an open-source collection of toolboxes (primarily written in MATLAB; with some components in C and Python) that aim to facilitate the acquisition and (computational) analysis of neuroimaging and behavioral data. Specifically, we reviewed the different toolboxes in TAPAS and highlighted how these might support the construction of end-to-end analysis pipelines – from raw data to clinical applications.

## 9 SOFTWARE NOTE

The **T**ranslational **A**lgorithms for **P**sychiatry-**A**dvancing **S**cience (TAPAS) software package, comprising all toolboxes described in this paper, is freely available as open-source code (https://www.translationalneuromodeling.org/tapas).

## ACKNOWLEDGEMENTS

This work was supported by the UZH Forschungskredit Postdoc, grant number FK-18-046, the ETH Zurich Postdoctoral Fellowship Program and the Marie Curie Actions for People COFUND Program, grant number FEL-49 15-2 (SF), a Marie Skłodowska-Curie Postdoctoral Fellowship from the European Union’s Horizon 2020 research and innovation program under the Grant Agreement No 793580, as well as by the Royal Society of New Zealand (OKH), the Strategic Focal Area “Personalized Health and Related Technologies (PHRT)” of the ETH Domain, grant number 2017-403 (SJH), as well as the René and Susanne Braginsky Foundation, the Clinical Research Priority Program “Pain” of the University of Zurich, and the Swiss National Science Foundation, grant number 320030_179377 (KES).

## Footnotes

1 ML is also used “on its own” in CP and applied directly to measured data, e.g., for producing patient-specific predictions or discovering structure in heterogeneous populations (Durstewitz et al., 2019; Dwyer et al., 2018; Gillan et al., 2017; Mourão-Miranda et al., 2012; Portugal et al., 2019; Schmaal et al., 2015). In this paper, however, we focus on approaches where ML operates on estimates provided by generative models.

2 Exteroception refers to perception of sensations originating from the external world, whereas interoception refers to perception of sensations originating from the own body or “internal world”.

3 Besides different variants of the HGF, the toolbox also implements various other behavioral models of learning, including the Rescorla-Wagner model (Rescorla and Wagner, 1972), Sutton model (Sutton, 1992), and a Hidden Markov model (Baum and Petrie, 1966).

## REFERENCES

Abraham, A., Pedregosa, F., Eickenberg, M., Gervais, P., Mueller, A., Kossaifi, J., Gramfort, A., Thirion, B., Varoquaux, G., 2014. Machine learning for neuroimaging with scikit-learn. Front Neuroinform 8, 14.

Adams, N.E., Hughes, L.E., Phillips, H.N., Shaw, A.D., Murley, A.G., Nesbitt, D., Cope, T.E., Bevan-Jones, W.R., Passamonti, L., Rowe, J.B., 2020. GABA-ergic Dynamics in Human Frontotemporal Networks Confirmed by Pharmaco-Magnetoencephalography. J Neurosci 40, 1640–1649.

Adams, R.A., Huys, Q.J., Roiser, J.P., 2016. Computational Psychiatry: towards a mathematically informed understanding of mental illness. J Neurol Neurosurg Psychiatry 87, 53–63.

Ahn, W.Y., Haines, N., Zhang, L., 2017. Revealing Neurocomputational Mechanisms of Reinforcement Learning and Decision-Making With the hBayesDM Package. Comput Psychiatr 1, 24–57.

Alfaro-Almagro, F., Jenkinson, M., Bangerter, N.K., Andersson, J.L.R., Griffanti, L., Douaud, G., Sotiropoulos, S.N., Jbabdi, S., Hernandez-Fernandez, M., Vallee, E., Vidaurre, D., Webster, M., McCarthy, P., Rorden, C., Daducci, A., Alexander, D.C., Zhang, H., Dragonu, I., Matthews, P.M., Miller, K.L., Smith, S.M., 2018. Image processing and Quality Control for the first 10,000 brain imaging datasets from UK Biobank. Neuroimage 166, 400–424.

Almeida, J.R., Versace, A., Mechelli, A., Hassel, S., Quevedo, K., Kupfer, D.J., Phillips, M.L., 2009. Abnormal amygdala-prefrontal effective connectivity to happy faces differentiates bipolar from major depression. Biol Psychiatry 66, 451–459.

Alsop, D.C., Detre, J.A., Golay, X., Gunther, M., Hendrikse, J., Hernandez-Garcia, L., Lu, H., MacIntosh, B.J., Parkes, L.M., Smits, M., van Osch, M.J., Wang, D.J., Wong, E.C., Zaharchuk, G., 2015. Recommended implementation of arterial spin-labeled perfusion MRI for clinical applications: A consensus of the ISMRM perfusion study group and the European consortium for ASL in dementia. Magn Reson Med 73, 102–116.

American Psychiatric Association, 2013. Diagnostic and Statistical Manual of Mental Disorders (DSM-5 R). American Psychiatric Publishing.

Aponte, E.A., Raman, S., Sengupta, B., Penny, W.D., Stephan, K.E., Heinzle, J., 2016. mpdcm: A toolbox for massively parallel dynamic causal modeling. J Neurosci Methods 257, 7–16.

Aponte, E.A., Schöbi, D., Stephan, K.E., Heinzle, J., 2017. The Stochastic Early Reaction, Inhibition, and late Action (SERIA) model for antisaccades. Plos Computational Biology 13.

Aponte, E.A., Schöbi, D., Stephan, K.E., Heinzle, J., 2020a. Computational Dissociation of Dopaminergic and Cholinergic Effects on Action Selection and Inhibitory Control. Biol Psychiatry Cogn Neurosci Neuroimaging 5, 364–372.

Aponte, E.A., Stephan, K.E., Heinzle, J., 2019. Switch costs in inhibitory control and voluntary behaviour: A computational study of the antisaccade task. Eur J Neurosci 50, 3205–3220.

Aponte, E.A., Tschan, D.G., Stephan, K.E., Heinzle, J., 2018. Inhibition failures and late errors in the antisaccade task: influence of cue delay. J Neurophysiol 120, 3001–3016.

Aponte, E.A., Yao, Y., Raman, S., Frässle, S., Heinzle, J., Penny, W.D., Stephan, K.E., 2020b. An introduction to thermodynamic integration and application to dynamic causal models.

Ashburner, J., 2016. Preparing fMRI Data for Statistical Analysis. Fmri Techniques and Protocols, 2nd Edition 119, 155–181.

Banerjee, S., Carlin, B.P., Gelfand, A.E., 2015. Hierarchical modeling and analysis for spatial data, Second edition. ed. CRC Press, Taylor & Francis Group, Boca Raton.

Baum, L.E., Petrie, T., 1966. Statistical Inference for Probabilistic Functions of Finite State Markov Chains. Annals of Mathematical Statistics 37, 1554-&.

Bellon, E.M., Haacke, E.M., Coleman, P.E., Sacco, D.C., Steiger, D.A., Gangarosa, R.E., 1986. MR artifacts: a review. AJR Am J Roentgenol 147, 1271–1281.

Birn, R.M., Diamond, J.B., Smith, M.A., Bandettini, P.A., 2006. Separating respiratory-variation-related fluctuations from neuronal-activity-related fluctuations in fMRI. Neuroimage 31, 1536–1548.

Birn, R.M., Smith, M.A., Jones, T.B., Bandettini, P.A., 2008. The respiration response function: the temporal dynamics of fMRI signal fluctuations related to changes in respiration. Neuroimage 40, 644–654.

Bishop, C.M., 2006. Pattern recognition and machine learning. Springer, New York. 12, 13, 47, 105.

Bollmann, S., Barth, M., 2020. New acquisition techniques and their prospects for the achievable resolution of fMRI. Prog Neurobiol, 101936.

Brazil, I.A., Mathys, C.D., Popma, A., Hoppenbrouwers, S.S., Cohn, M.D., 2017. Representational Uncertainty in the Brain During Threat Conditioning and the Link With Psychopathic Traits. Biol Psychiatry Cogn Neurosci Neuroimaging 2, 689–695.

Bright, M.G., Murphy, K., 2017. Cleaning up the fMRI time series: Mitigating noise with advanced acquisition and correction strategies. Neuroimage 154, 1–3.

Brodersen, K.H., Deserno, L., Schlagenhauf, F., Lin, Z., Penny, W.D., Buhmann, J.M., Stephan, K.E., 2014. Dissecting psychiatric spectrum disorders by generative embedding. Neuroimage Clin 4, 98–111.

Brodersen, K.H., Schofield, T.M., Leff, A.P., Ong, C.S., Lomakina, E.I., Buhmann, J.M., Stephan, K.E., 2011. Generative embedding for model-based classification of fMRI data. PLoS Comput Biol 7, e1002079.

Brooks, J.C., Beckmann, C.F., Miller, K.L., Wise, R.G., Porro, C.A., Tracey, I., Jenkinson, M., 2008. Physiological noise modelling for spinal functional magnetic resonance imaging studies. Neuroimage 39, 680–692.

Brosch, J.R., Talavage, T.M., Ulmer, J.L., Nyenhuis, J.A., 2002. Simulation of human respiration in fMRI with a mechanical model. IEEE Trans Biomed Eng 49, 700–707.

Browning, M., Carter, C.S., Chatham, C., Den Ouden, H., Gillan, C.M., Baker, J.T., Chekroud, A.M., Cools, R., Dayan, P., Gold, J., Goldstein, R.Z., Hartley, C.A., Kepecs, A., Lawson, R.P., Mourao-Miranda, J., Phillips, M.L., Pizzagalli, D.A., Powers, A., Rindskopf, D., Roiser, J.P., Schmack, K., Schiller, D., Sebold, M., Stephan, K.E., Frank, M.J., Huys, Q., Paulus, M., 2020. Realizing the Clinical Potential of Computational Psychiatry: Report From the Banbury Center Meeting, February 2019. Biol Psychiatry 88, e5–e10.

Bullmore, E.T., Frangou, S., Murray, R.M., 1997. The dysplastic net hypothesis: an integration of developmental and dysconnectivity theories of schizophrenia. Schizophr Res 28, 143–156.

Buxton, R., Wong, E., Frank, L., 1998. Dynamics of blood flow and oxygenation changes during brain activation: The balloon model. Magn. Reson. Med. 39, 855–864.

Calderhead, B., Girolami, M., 2009. Estimating Bayes factors via thermodynamic integration and population MCMC. Computational Statistics & Data Analysis 53, 4028–4045.

Carp, J., 2012. The secret lives of experiments: methods reporting in the fMRI literature. Neuroimage 63, 289–300.

Carter, C.S., Perlstein, W., Ganguli, R., Brar, J., Mintun, M., Cohen, J.D., 1998. Functional hypofrontality and working memory dysfunction in schizophrenia. Am J Psychiatry 155, 1285–1287.

Chang, C., Cunningham, J.P., Glover, G.H., 2009. Influence of heart rate on the BOLD signal: the cardiac response function. Neuroimage 44, 857–869.

Chang, C., Glover, G.H., 2009. Effects of model-based physiological noise correction on default mode network anti-correlations and correlations. Neuroimage 47, 1448–1459.

Cole, D.M., Diaconescu, A.O., Pfeiffer, U.J., Brodersen, K.H., Mathys, C.D., Julkowski, D., Ruhrmann, S., Schilbach, L., Tittgemeyer, M., Vogeley, K., Stephan, K.E., 2020. Atypical processing of uncertainty in individuals at risk for psychosis. Neuroimage Clin 26, 102239.

Courchesne, E., Pierce, K., Schumann, C.M., Redcay, E., Buckwalter, J.A., Kennedy, D.P., Morgan, J., 2007. Mapping early brain development in autism. Neuron 56, 399–413.

Cox, R.W., 1996. AFNI: software for analysis and visualization of functional magnetic resonance neuroimages. Comput Biomed Res 29, 162–173.

Curtis, C.E., Calkins, M.E., Grove, W.M., Feil, K.J., Lacono, W.G., 2001. Saccadic disinhibition in patients with acute and remitted schizophrenia and their first-degree biological relatives. American Journal of Psychiatry 158, 100–106.

Cuthbert, B., Insel, T., 2010. Toward New Approaches to Psychotic Disorders: The NIMH Research Domain Criteria Project. Schizophrenia Bull 36, 1061–1062.

Dagli, M.S., Ingeholm, J.E., Haxby, J.V., 1999. Localization of cardiac-induced signal change in fMRI. Neuroimage 9, 407–415.

Daunizeau, J., Adam, V., Rigoux, L., 2014. VBA: a probabilistic treatment of nonlinear models for neurobiological and behavioural data. PLoS Comput Biol 10, e1003441.

Daunizeau, J., David, O., Stephan, K., 2011. Dynamic causal modelling: A critical review of the biophysical and statistical foundations. Neuroimage 58, 312–322.

Daunizeau, J., den Ouden, H.E., Pessiglione, M., Kiebel, S.J., Friston, K.J., Stephan, K.E., 2010a. Observing the observer (II): deciding when to decide. Plos One 5, e15555.

Daunizeau, J., den Ouden, H.E., Pessiglione, M., Kiebel, S.J., Stephan, K.E., Friston, K.J., 2010b. Observing the observer (I): meta-bayesian models of learning and decision-making. Plos One 5, e15554.

David, O., Kiebel, S.J., Harrison, L.M., Mattout, J., Kilner, J.M., Friston, K.J., 2006. Dynamic causal modeling of evoked responses in EEG and MEG. Neuroimage 30, 1255–1272.

Deco, G., Kringelbach, M.L., 2014. Great expectations: using whole-brain computational connectomics for understanding neuropsychiatric disorders. Neuron 84, 892–905.

Deserno, L., Sterzer, P., Wüstenberg, T., Heinz, A., Schlagenhauf, F., 2012. Reduced prefrontal-parietal effective connectivity and working memory deficits in schizophrenia. J Neurosci 32, 12–20.

Di Martino, A., Yan, C.G., Li, Q., Denio, E., Castellanos, F.X., Alaerts, K., Anderson, J.S., Assaf, M., Bookheimer, S.Y., Dapretto, M., Deen, B., Delmonte, S., Dinstein, I., Ertl-Wagner, B., Fair, D.A., Gallagher, L., Kennedy, D.P., Keown, C.L., Keysers, C., Lainhart, J.E., Lord, C., Luna, B., Menon, V., Minshew, N.J., Monk, C.S., Mueller, S., Muller, R.A., Nebel, M.B., Nigg, J.T., O’Hearn, K., Pelphrey, K.A., Peltier, S.J., Rudie, J.D., Sunaert, S., Thioux, M., Tyszka, J.M., Uddin, L.Q., Verhoeven, J.S., Wenderoth, N., Wiggins, J.L., Mostofsky, S.H., Milham, M.P., 2014. The autism brain imaging data exchange: towards a large-scale evaluation of the intrinsic brain architecture in autism. Mol Psychiatry 19, 659–667.

Dima, D., Roiser, J.P., Dietrich, D.E., Bonnemann, C., Lanfermann, H., Emrich, H.M., Dillo, W., 2009. Understanding why patients with schizophrenia do not perceive the hollow-mask illusion using dynamic causal modelling. Neuroimage 46, 1180–1186.

Durstewitz, D., Koppe, G., Meyer-Lindenberg, A., 2019. Deep neural networks in psychiatry. Mol Psychiatry 24, 1583–1598.

Dwyer, D.B., Falkai, P., Koutsouleris, N., 2018. Machine Learning Approaches for Clinical Psychology and Psychiatry. Annu Rev Clin Psychol 14, 91–118.

Dymerska, B., Poser, B.A., Barth, M., Trattnig, S., Robinson, S.D., 2018. A method for the dynamic correction of B0-related distortions in single-echo EPI at 7T. Neuroimage 168, 321–331.

Efron, B., Morris, C., 1973. Stein’s estimation rule and its competitors - Empirical Bayes Approach. J Am Stat Assoc 68, 117–130.

Esteban, O., Birman, D., Schaer, M., Koyejo, O.O., Poldrack, R.A., Gorgolewski, K.J., 2017. MRIQC: Advancing the automatic prediction of image quality in MRI from unseen sites. Plos One 12, e0184661.

Esteban, O., Markiewicz, C.J., Blair, R.W., Moodie, C.A., Isik, A.I., Erramuzpe, A., Kent, J.D., Goncalves, M., DuPre, E., Snyder, M., Oya, H., Ghosh, S.S., Wright, J., Durnez, J., Poldrack, R.A., Gorgolewski, K.J., 2019. fMRIPrep: a robust preprocessing pipeline for functional MRI. Nat Methods 16, 111–116.

Filiou, M.D., Turck, C.W., 2011. General overview: biomarkers in neuroscience research. Int Rev Neurobiol 101, 1–17.

Fischl, B., 2012. FreeSurfer. Neuroimage 62, 774–781.

Frässle, S., Harrison, S.J., Heinzle, J., Clementz, B., Tamminga, C., Sweeney, J., Gershon, E.S., Keshavan, M., Pearlson, G., Powers, A., Stephan, K.E., 2020a. Regression dynamic causal modeling for resting-state fMRI. doi: https://doi.org/10.1101/2020.08.12.247536.

Frässle, S., Lomakina, E.I., Kasper, L., Manjaly, Z.M., Leff, A., Pruessmann, K.P., Buhmann, J.M., Stephan, K.E., 2018a. A generative model of whole-brain effective connectivity. Neuroimage 179, 505–529.

Frässle, S., Lomakina, E.I., Razi, A., Friston, K.J., Buhmann, J.M., Stephan, K.E., 2017. Regression DCM for fMRI. Neuroimage 155, 406–421.

Frässle, S., Manjaly, Z.M., Do, C.T., Kasper, L., Pruessmann, K.P., Stephan, K.E., 2021. Whole-brain estimates of directed connectivity for human connectomics. Neuroimage 225, 117491.

Frässle, S., Marquand, A.F., Schmaal, L., Dinga, R., Veltman, D.J., van der Wee, N.J.A., van Tol, M.J., Schöbi, D., Penninx, B., Stephan, K.E., 2020b. Predicting individual clinical trajectories of depression with generative embedding. Neuroimage Clin 26, 102213.

Frässle, S., Stephan, K.E., Friston, K.J., Steup, M., Krach, S., Paulus, F.M., Jansen, A., 2015. Test-retest reliability of dynamic causal modeling for fMRI. Neuroimage 117, 56–66.

Frässle, S., Yao, Y., Schöbi, D., Aponte, E.A., Heinzle, J., Stephan, K.E., 2018b. Generative models for clinical applications in computational psychiatry. Wiley Interdiscip Rev Cogn Sci 9, e1460.

Friston, K., 2005a. A theory of cortical responses. Philos Trans R Soc Lond B Biol Sci 360, 815–836.

Friston, K., 2009. The free-energy principle: a rough guide to the brain? Trends Cogn Sci 13, 293–301.

Friston, K., Brown, H.R., Siemerkus, J., Stephan, K.E., 2016a. The dysconnection hypothesis (2016). Schizophr Res 176, 83–94.

Friston, K., Harrison, L., Penny, W., 2003. Dynamic causal modelling. Neuroimage 19, 1273–1302.

Friston, K., Mattout, J., Trujillo-Barreto, N., Ashburner, J., Penny, W., 2007. Variational free energy and the Laplace approximation. Neuroimage 34, 220–234.

Friston, K., Moran, R., Seth, A., 2013. Analysing connectivity with Granger causality and dynamic causal modelling. Current Opinion in Neurobiology 23, 172–178.

Friston, K.J., 2005b. Hallucinations and perceptual inference. Behavioral and Brain Sciences 28, 764-+.

Friston, K.J., 2011. Functional and effective connectivity: a review. Brain Connect 1, 13–36.

Friston, K.J., Ashburner, J.T., Kiebel, S.J., Nichols, T.E., Penny, W.D., 2006. Statistical Parametric Mapping : the Analysis of Functional Brain Images. Elsevier, Burlington.

Friston, K.J., Frith, C.D., 1995. Schizophrenia: a disconnection syndrome? Clin Neurosci 3, 89–97.

Friston, K.J., Litvak, V., Oswal, A., Razi, A., Stephan, K.E., van Wijk, B.C., Ziegler, G., Zeidman, P., 2016b. Bayesian model reduction and empirical Bayes for group (DCM) studies. Neuroimage 128, 413–431.

Friston, K.J., Mechelli, A., Turner, R., Price, C.J., 2000. Nonlinear responses in fMRI: the Balloon model, Volterra kernels, and other hemodynamics. Neuroimage 12, 466–477.

Friston, K.J., Redish, A.D., Gordon, J.A., 2017. Computational Nosology and Precision Psychiatry. Comput Psychiatr 1, 2–23.

Friston, K.J., Stephan, K.E., Montague, R., Dolan, R.J., 2014. Computational psychiatry: the brain as a phantastic organ. Lancet Psychiatry 1, 148–158.

Friston, K.J., Williams, S., Howard, R., Frackowiak, R.S., Turner, R., 1996. Movement-related effects in fMRI time-series. Magn Reson Med 35, 346–355.

Fukushima, J., Fukushima, K., Chiba, T., Tanaka, S., Yamashita, I., Kato, M., 1988. Disturbances of Voluntary Control of Saccadic Eye-Movements in Schizophrenic-Patients. Biological Psychiatry 23, 670–677.

Gardner, E.A., Ellis, J.H., Hyde, R.J., Aisen, A.M., Quint, D.J., Carson, P.L., 1995. Detection of degradation of magnetic resonance (MR) images: comparison of an automated MR image-quality analysis system with trained human observers. Acad Radiol 2, 277–281.

Gedamu, E.L., Collins, D.L., Arnold, D.L., 2008. Automated quality control of brain MR images. J Magn Reson Imaging 28, 308–319.

Gelman, A., Charlin, J.B., Stern, H.S., Rubin, D.B., 2004. Bayesian Data Analysis. Chapman and Hall.

Gilbert, J.R., Symmonds, M., Hanna, M.G., Dolan, R.J., Friston, K.J., Moran, R.J., 2016. Profiling neuronal ion channelopathies with non-invasive brain imaging and dynamic causal models: Case studies of single gene mutations. Neuroimage 124, 43–53.

Gillan, C.M., Whelan, R., 2017. What big data can do for treatment in psychiatry. Current Opinion in Behavioral Sciences 18, 34–42.

Glasser, M.F., Sotiropoulos, S.N., Wilson, J.A., Coalson, T.S., Fischl, B., Andersson, J.L., Xu, J., Jbabdi, S., Webster, M., Polimeni, J.R., Van Essen, D.C., Jenkinson, M., Consortium, W.U.-M.H., 2013. The minimal preprocessing pipelines for the Human Connectome Project. Neuroimage 80, 105–124.

Glover, G.H., Li, T.Q., Ress, D., 2000. Image-based method for retrospective correction of physiological motion effects in fMRI: RETROICOR. Magn Reson Med 44, 162–167.

Gorgolewski, K.J., Auer, T., Calhoun, V.D., Craddock, R.C., Das, S., Duff, E.P., Flandin, G., Ghosh, S.S., Glatard, T., Halchenko, Y.O., Handwerker, D.A., Hanke, M., Keator, D., Li, X., Michael, Z., Maumet, C., Nichols, B.N., Nichols, T.E., Pellman, J., Poline, J.B., Rokem, A., Schaefer, G., Sochat, V., Triplett, W., Turner, J.A., Varoquaux, G., Poldrack, R.A., 2016. The brain imaging data structure, a format for organizing and describing outputs of neuroimaging experiments. Sci Data 3, 160044.

Greicius, M.D., Flores, B.H., Menon, V., Glover, G.H., Solvason, H.B., Kenna, H., Reiss, A.L., Schatzberg, A.F., 2007. Resting-state functional connectivity in major depression: abnormally increased contributions from subgenual cingulate cortex and thalamus. Biol Psychiatry 62, 429–437.

Grèzes, J., Wicker, B., Berthoz, S., de Gelder, B., 2009. A failure to grasp the affective meaning of actions in autism spectrum disorder subjects. Neuropsychologia 47, 1816–1825.

Hallett, P.E., 1978. Primary and secondary saccades to goals defined by instructions. Vision Res 18, 1279–1296.

Harrison, O.K., Garfinkel, S.N., Marlow, L., Finnegan, S., Marino, S., Nanz, L., Allen, M., Finnemann, J., Keur-Huizinga, L., Harrison, S.J., Stephan, K.E., Pattinson, K., Fleming, S.M., 2020a. The Filter Detection Task for measurement of breathing-related interoception and metacognition.

Harrison, S.J., Bianchi, S., Heinzle, J., Stephan, K.E., Iglesias, S., Kasper, L., 2020b. A Hilbert-based method for processing respiratory timeseries. https://doi.org/10.1101/2020.09.30.321562.

Havlicek, M., Roebroeck, A., Friston, K., Gardumi, A., Ivanov, D., Uludag, K., 2015. Physiologically informed dynamic causal modeling of fMRI data. Neuroimage 122, 355–372.

Havlicek, M., Uludag, K., 2020. A dynamical model of the laminar BOLD response. Neuroimage 204, 116209.

Heinzle, J., Aponte, E.A., Stephan, K.E., 2016a. Computational models of eye movements and their application to schizophrenia. Current Opinion in Behavioral Sciences 11, 21–29.

Heinzle, J., Koopmans, P.J., den Ouden, H.E.M., Raman, S., Stephan, K.E., 2016b. A hemodynamic model for layered BOLD signals. Neuroimage 125, 556–570.

Helske, J., 2017. KFAS: Exponential Family State Space Models in R. Journal of Statistical Software 78, 1–39.

Hendriks, A.D., D’Agata, F., Raimondo, L., Schakel, T., Geerts, L., Luijten, P.R., Klomp, D.W.J., Petridou, N., 2020. Pushing functional MRI spatial and temporal resolution further: High-density receive arrays combined with shot-selective 2D CAIPIRINHA for 3D echo-planar imaging at 7 T. NMR Biomed 33, e4281.

Hesselmann, V., Girnus, R., Wedekind, C., Hunsche, S., Bunke, J., Schulte, O., Sorger, B., Lasek, K., Krug, B., Sturm, V., Lackner, K., 2004. Functional MRI using multiple receiver coils: BOLD signal changes and signal-to-noise ratio for three-dimensional-PRESTO vs. single shot EPI in comparison to a standard quadrature head coil. J Magn Reson Imaging 20, 321–326.

Hirsch, J.A., Bishop, B., 1981. Respiratory Sinus Arrhythmia in Humans -How Breathing Pattern Modulates Heart-Rate. American Journal of Physiology 241, H620–H629.

Huber, L., Handwerker, D.A., Jangraw, D.C., Chen, G., Hall, A., Stuber, C., Gonzalez-Castillo, J., Ivanov, D., Marrett, S., Guidi, M., Goense, J., Poser, B.A., Bandettini, P.A., 2017. High-Resolution CBV-fMRI Allows Mapping of Laminar Activity and Connectivity of Cortical Input and Output in Human M1. Neuron 96, 1253–1263 e1257.

Huettel, S.A., Song, A.W., McCarthy, G., 2009. Functional magnetic resonance imaging. Sinauer Associates, Sunderland, Mass.

Hutton, C., Josephs, O., Stadler, J., Featherstone, E., Reid, A., Speck, O., Bernarding, J., Weiskopf, N., 2011. The impact of physiological noise correction on fMRI at 7 T. Neuroimage 57, 101–112.

Hutton, S.B., Ettinger, U., 2006. The antisaccade task as a research tool in psychopathology: A critical review. Psychophysiology 43, 302–313.

Huys, Q.J., Maia, T.V., Frank, M.J., 2016. Computational psychiatry as a bridge from neuroscience to clinical applications. Nat Neurosci 19, 404–413.

Iglesias, S., Kasper, L., Harrison, S.J., Manka, R., Mathys, C., Stephan, K.E., 2020. Cholinergic and dopaminergic effects on prediction error and uncertainty responses during sensory associative learning. Neuroimage, 117590.

Iglesias, S., Mathys, C., Brodersen, K.H., Kasper, L., Piccirelli, M., den Ouden, H.E., Stephan, K.E., 2013. Hierarchical prediction errors in midbrain and basal forebrain during sensory learning. Neuron 80, 519–530.

Iglesias, S., Tomiello, S., Schneebeli, M., Stephan, K.E., 2016. Models of neuromodulation for computational psychiatry. Wiley Interdiscip Rev Cogn Sci.

Jenkinson, M., Beckmann, C.F., Behrens, T.E., Woolrich, M.W., Smith, S.M., 2012. Fsl. Neuroimage 62, 782–790.

Jezzard, P., Balaban, R.S., 1995. Correction for geometric distortion in echo planar images from B0 field variations. Magn Reson Med 34, 65–73.

Jezzard, P., Clare, S., 1999. Sources of distortion in functional MRI data. Hum Brain Mapp 8, 80–85.

Jirsa, V.K., Proix, T., Perdikis, D., Woodman, M.M., Wang, H., Gonzalez-Martinez, J., Bernard, C., Bénar, C., Guye, M., Chauvel, P., Bartolomei, F., 2016. The Virtual Epileptic Patient: Individualized whole-brain models of epilepsy spread. Neuroimage.

Jirsa, V.K., Sporns, O., Breakspear, M., Deco, G., McIntosh, A.R., 2010. Towards the virtual brain: network modeling of the intact and the damaged brain. Arch Ital Biol 148, 189–205.

Kapur, S., Phillips, A.G., Insel, T.R., 2012. Why has it taken so long for biological psychiatry to develop clinical tests and what to do about it? Mol Psychiatry 17, 1174–1179.

Kasper, L., Bollmann, S., Diaconescu, A.O., Hutton, C., Heinzle, J., Iglesias, S., Hauser, T.U., Sebold, M., Manjaly, Z.M., Pruessmann, K.P., Stephan, K.E., 2017. The PhysIO Toolbox for Modeling Physiological Noise in fMRI Data. J Neurosci Methods 276, 56–72.

Kass, R., Steffey, D., 1989. Aproximate Bayesian inference in conditionally indepedent hierarchical models (parametric empirical Bayes models). J Am Stat Assoc 84, 717–726.

Keil, B., Wald, L.L., 2013. Massively parallel MRI detector arrays. Journal of Magnetic Resonance 229, 75–89.

Kennedy, D.P., Redcay, E., Courchesne, E., 2006. Failing to deactivate: resting functional abnormalities in autism. Proc Natl Acad Sci U S A 103, 8275–8280.

Khalsa, S.S., Adolphs, R., Cameron, O.G., Critchley, H.D., Davenport, P.W., Feinstein, J.S., Feusner, J.D., Garfinkel, S.N., Lane, R.D., Mehling, W.E., Meuret, A.E., Nemeroff, C.B., Oppenheimer, S., Petzschner, F.H., Pollatos, O., Rhudy, J.L., Schramm, L.P., Simmons, W.K., Stein, M.B., Stephan, K.E., Van den Bergh, O., Van Diest, I., von Leupoldt, A., Paulus, M.P., Interoception Summit, p., 2018. Interoception and Mental Health: A Roadmap. Biol Psychiatry Cogn Neurosci Neuroimaging 3, 501–513.

Kirilina, E., Lutti, A., Poser, B.A., Blankenburg, F., Weiskopf, N., 2016. The quest for the best: The impact of different EPI sequences on the sensitivity of random effect fMRI group analyses. Neuroimage 126, 49–59.

Kirkwood, J.G., 1935. Statistical mechanics of fluid mixtures. Journal of Chemical Physics 3, 300–313.

Koretsky, A.P., 2012. Early development of arterial spin labeling to measure regional brain blood flow by MRI. Neuroimage 62, 602–607.

Kruger, G., Glover, G.H., 2001. Physiological noise in oxygenation-sensitive magnetic resonance imaging. Magn Reson Med 46, 631–637.

Krystal, J.H., State, M.W., 2014. Psychiatric disorders: diagnosis to therapy. Cell 157, 201–214.

Kundu, P., Voon, V., Balchandani, P., Lombardo, M.V., Poser, B.A., Bandettini, P.A., 2017. Multi-echo fMRI: A review of applications in fMRI denoising and analysis of BOLD signals. Neuroimage 154, 59–80.

Lartillot, N., Philippe, H., 2006. Computing Bayes factors using thermodynamic integration. Systematic Biology 55, 195–207.

Lawson, R.P., Mathys, C., Rees, G., 2017. Adults with autism overestimate the volatility of the sensory environment. Nat Neurosci 20, 1293–1299.

Lefebvre, S., Demeulemeester, M., Leroy, A., Delmaire, C., Lopes, R., Pins, D., Thomas, P., Jardri, R., 2016. Network dynamics during the different stages of hallucinations in schizophrenia. Hum Brain Mapp 37, 2571–2586.

Li, B., Cui, L.B., Xi, Y.B., Friston, K.J., Guo, F., Wang, H.N., Zhang, L.C., Bai, Y.H., Tan, Q.R., Yin, H., Lu, H., 2017. Abnormal Effective Connectivity in the Brain is Involved in Auditory Verbal Hallucinations in Schizophrenia. Neurosci Bull 33, 281–291.

Lu, H., Hua, J., van Zijl, P.C., 2013. Noninvasive functional imaging of cerebral blood volume with vascular-space-occupancy (VASO) MRI. NMR Biomed 26, 932–948.

Maia, T., Frank, M., 2011. From reinforcement learning models to psychiatric and neurological disorders. Nat Neurosci 14, 154–162.

Maritz, J.S., Lwin, T., 1989. Empirical Bayes methods, 2nd ed. Chapman and Hall, London ; New York.

Marshall, L., Mathys, C., Ruge, D., de Berker, A.O., Dayan, P., Stephan, K.E., Bestmann, S., 2016. Pharmacological Fingerprints of Contextual Uncertainty. PLoS Biol 14, e1002575.

Mathys, C., Daunizeau, J., Friston, K.J., Stephan, K.E., 2011. A bayesian foundation for individual learning under uncertainty. Front Hum Neurosci 5, 39.

Mathys, C.D., Lomakina, E.I., Daunizeau, J., Iglesias, S., Brodersen, K.H., Friston, K.J., Stephan, K.E., 2014. Uncertainty in perception and the Hierarchical Gaussian Filter. Front Hum Neurosci 8, 825.

Maxwell, S.E., 2004. The persistence of underpowered studies in psychological research: causes, consequences, and remedies. Psychol Methods 9, 147–163.

Mayberg, H.S., 1997. Limbic-cortical dysregulation: a proposed model of depression. J Neuropsychiatry Clin Neurosci 9, 471–481.

Menon, R.S., 2002. Postacquisition suppression of large-vessel BOLD signals in high-resolution fMRI. Magn Reson Med 47, 1–9.

Menon, V., 2011. Large-scale brain networks and psychopathology: a unifying triple network model. Trends Cogn Sci 15, 483–506.

Montague, P., Dolan, R., Friston, K., Dayan, P., 2012. Computational psychiatry. Trends Cogn Sci 16, 72–80.

Moran, R., Pinotsis, D.A., Friston, K., 2013. Neural masses and fields in dynamic causal modeling. Front Comput Neurosci 7, 57.

Moran, R.J., Jung, F., Kumagai, T., Endepols, H., Graf, R., Dolan, R.J., Friston, K.J., Stephan, K.E., Tittgemeyer, M., 2011. Dynamic causal models and physiological inference: a validation study using isoflurane anaesthesia in rodents. Plos One 6, e22790.

Mortamet, B., Bernstein, M.A., Jack, C.R., Jr., Gunter, J.L., Ward, C., Britson, P.J., Meuli, R., Thiran, J.P., Krueger, G., Alzheimer’s Disease Neuroimaging, I., 2009. Automatic quality assessment in structural brain magnetic resonance imaging. Magn Reson Med 62, 365–372.

Mourao-Miranda, J., Reinders, A.A.T.S., Rocha-Rego, V., Lappin, J., Rondina, J., Morgan, C., Morgan, K.D., Fearon, P., Jones, P.B., Doody, G.A., Murray, R.M., Kapur, S., Dazzan, P., 2012. Individualized prediction of illness course at the first psychotic episode: a support vector machine MRI study. Psychological Medicine 42, 1037–1047.

Muller, R.A., 2007. The study of autism as a distributed disorder. Ment Retard Dev Disabil Res Rev 13, 85–95.

Murphy, K., Birn, R.M., Bandettini, P.A., 2013. Resting-state fMRI confounds and cleanup. Neuroimage 80, 349–359.

Owen, M.J., 2014. New approaches to psychiatric diagnostic classification. Neuron 84, 564–571.

Paulus, M.P., Huys, Q.J., Maia, T.V., 2016. A Roadmap for the Development of Applied Computational Psychiatry. Biol Psychiatry Cogn Neurosci Neuroimaging 1, 386–392.

Pedregosa, F., Varoquaux, G., Gramfort, A., Michel, V., Thirion, B., Grisel, O., Blondel, M., Prettenhofer, P., Weiss, R., Dubourg, V., Vanderplas, J., Passos, A., Cournapeau, D., Brucher, M., Perrot, M., Duchesnay, E., 2011. Scikit-learn: Machine Learning in Python. Journal of Machine Learning Research 12, 2825–2830.

Pellicano, E., Burr, D., 2012. When the world becomes ’too real’: a Bayesian explanation of autistic perception. Trends in Cognitive Sciences 16, 504–510.

Penny, W.D., Stephan, K.E., Mechelli, A., Friston, K.J., 2004. Comparing dynamic causal models. Neuroimage 22, 1157–1172.

Petzschner, F.H., Weber, L.A., Wellstein, K.V., Paolini, G., Do, C.T., Stephan, K.E., 2019. Focus of attention modulates the heartbeat evoked potential. Neuroimage 186, 595–606.

Petzschner, F.H., Weber, L.A.E., Gard, T., Stephan, K.E., 2017. Computational Psychosomatics and Computational Psychiatry: Toward a Joint Framework for Differential Diagnosis. Biol Psychiatry.

Pizarro, R.A., Cheng, X., Barnett, A., Lemaitre, H., Verchinski, B.A., Goldman, A.L., Xiao, E., Luo, Q., Berman, K.F., Callicott, J.H., Weinberger, D.R., Mattay, V.S., 2016. Automated Quality Assessment of Structural Magnetic Resonance Brain Images Based on a Supervised Machine Learning Algorithm. Front Neuroinform 10, 52.

Poldrack, R.A., Congdon, E., Triplett, W., Gorgolewski, K.J., Karlsgodt, K.H., Mumford, J.A., Sabb, F.W., Freimer, N.B., London, E.D., Cannon, T.D., Bilder, R.M., 2016. A phenome-wide examination of neural and cognitive function. Sci Data 3, 160110.

Poldrack, R.A., Gorgolewski, K.J., 2014. Making big data open: data sharing in neuroimaging. Nat Neurosci 17, 1510–1517.

Poline, J.B., Breeze, J.L., Ghosh, S., Gorgolewski, K., Halchenko, Y.O., Hanke, M., Haselgrove, C., Helmer, K.G., Keator, D.B., Marcus, D.S., Poldrack, R.A., Schwartz, Y., Ashburner, J., Kennedy, D.N., 2012. Data sharing in neuroimaging research. Front Neuroinform 6, 9.

Portugal, L.C.L., Schrouff, J., Stiffler, R., Bertocci, M., Bebko, G., Chase, H., Lockovitch, J., Aslam, H., Graur, S., Greenberg, T., Pereira, M., Oliveira, L., Phillips, M., Mourao-Miranda, J., 2019. Predicting anxiety from wholebrain activity patterns to emotional faces in young adults: a machine learning approach. Neuroimage-Clinical 23.

Poser, B.A., Setsompop, K., 2018. Pulse sequences and parallel imaging for high spatiotemporal resolution MRI at ultra-high field. Neuroimage 168, 101–118.

Poser, B.A., Versluis, M.J., Hoogduin, J.M., Norris, D.G., 2006. BOLD contrast sensitivity enhancement and artifact reduction with multiecho EPI: parallel-acquired inhomogeneity-desensitized fMRI. Magn Reson Med 55, 1227–1235.

Posse, S., Wiese, S., Gembris, D., Mathiak, K., Kessler, C., Grosse-Ruyken, M.L., Elghahwagi, B., Richards, T., Dager, S.R., Kiselev, V.G., 1999. Enhancement of BOLD-contrast sensitivity by single-shot multi-echo functional MR imaging. Magn Reson Med 42, 87–97.

Power, J.D., Barnes, K.A., Snyder, A.Z., Schlaggar, B.L., Petersen, S.E., 2012. Spurious but systematic correlations in functional connectivity MRI networks arise from subject motion. Neuroimage 59, 2142–2154.

Power, J.D., Mitra, A., Laumann, T.O., Snyder, A.Z., Schlaggar, B.L., Petersen, S.E., 2014. Methods to detect, characterize, and remove motion artifact in resting state fMRI. Neuroimage 84, 320–341.

Power, J.D., Plitt, M., Laumann, T.O., Martin, A., 2017. Sources and implications of whole-brain fMRI signals in humans. Neuroimage 146, 609–625.

Powers, A.R., Mathys, C., Corlett, P.R., 2017. Pavlovian conditioning-induced hallucinations result from overweighting of perceptual priors. Science 357, 596–600.

Radulescu, E., Minati, L., Ganeshan, B., Harrison, N.A., Gray, M.A., Beacher, F.D., Chatwin, C., Young, R.C., Critchley, H.D., 2013. Abnormalities in fronto-striatal connectivity within language networks relate to differences in grey-matter heterogeneity in Asperger syndrome. Neuroimage Clin 2, 716–726.

Raj, D., Anderson, A.W., Gore, J.C., 2001. Respiratory effects in human functional magnetic resonance imaging due to bulk susceptibility changes. Physics in Medicine and Biology 46, 3331–3340.

Raman, S., Deserno, L., Schlagenhauf, F., Stephan, K.E., 2016. A hierarchical model for integrating unsupervised generative embedding and empirical Bayes. J Neurosci Methods 269, 6–20.

Rao, R.P., Ballard, D.H., 1999. Predictive coding in the visual cortex: a functional interpretation of some extra-classical receptive-field effects. Nat Neurosci 2, 79–87.

Reilly, J.L., Frankovich, K., Hill, S., Gershon, E.S., Keefe, R.S.E., Keshavan, M.S., Pearlson, G.D., Tamminga, C.A., Sweeney, J.A., 2014. Elevated Antisaccade Error Rate as an Intermediate Phenotype for Psychosis Across Diagnostic Categories. Schizophrenia Bulletin 40, 1011–1021.

Rescorla, R.A., Wagner, A.R., 1972. A theory of Pavlovian conditioning: variations in the effectiveness of reinforcement and nonreinforcement. In: Black AH, Prokasy WF, editors. Classical conditioning II: current research and theory. New York: Appleton Century Crofts., 64–99.

Reuter, M., Tisdall, M.D., Qureshi, A., Buckner, R.L., van der Kouwe, A.J.W., Fischl, B., 2015. Head motion during MRI acquisition reduces gray matter volume and thickness estimates. Neuroimage 107, 107–115.

Rieger, S.W., Stephan, K.E., Harrison, O.K., 2020. Remote, Automated, and MRI-Compatible Administration of Interoceptive Inspiratory Resistive Loading. Front Hum Neurosci 14, 161.

Rommelse, N.N.J., Van der Stigchel, S., Sergeant, J.A., 2008. A review on eye movement studies in childhood and adolescent psychiatry. Brain and Cognition 68, 391–414.

Rowe, D.B., 2005. Parameter estimation in the magnitude-only and complex-valued fMRI data models. Neuroimage 25, 1124–1132.

Schlösser, R.G., Wagner, G., Koch, K., Dahnke, R., Reichenbach, J.R., Sauer, H., 2008. Fronto-cingulate effective connectivity in major depression: a study with fMRI and dynamic causal modeling. Neuroimage 43, 645–655.

Schmaal, L., Marquand, A.F., Rhebergen, D., van Tol, M.J., Ruhé, H.G., van der Wee, N.J., Veltman, D.J., Penninx, B.W., 2015. Predicting the Naturalistic Course of Major Depressive Disorder Using Clinical and Multimodal Neuroimaging Information: A Multivariate Pattern Recognition Study. Biol Psychiatry 78, 278–286.

Schofield, T.M., Penny, W.D., Stephan, K.E., Crinion, J.T., Thompson, A.J., Price, C.J., Leff, A.P., 2012. Changes in auditory feedback connections determine the severity of speech processing deficits after stroke. J Neurosci 32, 4260–4270.

Sengupta, B., Friston, K.J., Penny, W.D., 2015. Gradient-free MCMC methods for dynamic causal modelling. Neuroimage 112, 375–381.

Sengupta, B., Friston, K.J., Penny, W.D., 2016. Gradient-based MCMC samplers for dynamic causal modelling. Neuroimage 125, 1107–1118.

Shawe-Taylor, J., Cristianini, N., 2004. Kernel Methods for Pattern Analysis. cambridge University Press.

Smith, S.M., Jenkinson, M., Woolrich, M.W., Beckmann, C.F., Behrens, T.E.J., Johansen-Berg, H., Bannister, P.R., De Luca, M., Drobnjak, I., Flitney, D.E., Niazy, R.K., Saunders, J., Vickers, J., Zhang, Y.Y., De Stefano, N., Brady, J.M., Matthews, P.M., 2004. Advances in functional and structural MR image analysis and implementation as FSL. Neuroimage 23, S208–S219.

Snoek, L., Miletic, S., Scholte, H.S., 2019. How to control for confounds in decoding analyses of neuroimaging data. Neuroimage 184, 741–760.

Stalnaker, T.A., Howard, J.D., Takahashi, Y.K., Gershman, S.J., Kahnt, T., Schoenbaum, G., 2019. Dopamine neuron ensembles signal the content of sensory prediction errors. Elife 8.

Stephan, K., Baldeweg, T., Friston, K., 2006. Synaptic plasticity and dysconnection in schizophrenia. Biol Psychiatry 59, 929–939.

Stephan, K., Friston, K., Frith, C., 2009. Dysconnection in Schizophrenia: From Abnormal Synaptic Plasticity to Failures of Self-monitoring. Schizophrenia Bulletin 35, 509–527.

Stephan, K., Mathys, C., 2014. Computational approaches to psychiatry. Curr Opin Neurobiol 25, 85–92.

Stephan, K., Penny, W., Moran, R., den Ouden, H., Daunizeau, J., Friston, K., 2010. Ten simple rules for dynamic causal modeling. Neuroimage 49, 3099–3109.

Stephan, K.E., Bach, D.R., Fletcher, P.C., Flint, J., Frank, M.J., Friston, K.J., Heinz, A., Huys, Q.J., Owen, M.J., Binder, E.B., Dayan, P., Johnstone, E.C., Meyer-Lindenberg, A., Montague, P.R., Schnyder, U., Wang, X.J., Breakspear, M., 2016a. Charting the landscape of priority problems in psychiatry, part 1: classification and diagnosis. Lancet Psychiatry 3, 77–83.

Stephan, K.E., Binder, E.B., Breakspear, M., Dayan, P., Johnstone, E.C., Meyer-Lindenberg, A., Schnyder, U., Wang, X.J., Bach, D.R., Fletcher, P.C., Flint, J., Frank, M.J., Heinz, A., Huys, Q.J., Montague, P.R., Owen, M.J., Friston, K.J., 2016b. Charting the landscape of priority problems in psychiatry, part 2: pathogenesis and aetiology. Lancet Psychiatry 3, 84–90.

Stephan, K.E., Iglesias, S., Heinzle, J., Diaconescu, A.O., 2015. Translational Perspectives for Computational Neuroimaging. Neuron 87, 716–732.

Stephan, K.E., Schlagenhauf, F., Huys, Q.J., Raman, S., Aponte, E.A., Brodersen, K.H., Rigoux, L., Moran, R.J., Daunizeau, J., Dolan, R.J., Friston, K.J., Heinz, A., 2017. Computational neuroimaging strategies for single patient predictions. Neuroimage 145, 180–199.

Stephan, K.E., Weiskopf, N., Drysdale, P.M., Robinson, P.A., Friston, K.J., 2007. Comparing hemodynamic models with DCM. Neuroimage 38, 387–401.

Strother, S.C., 2006. Evaluating fMRI preprocessing pipelines. IEEE Eng Med Biol Mag 25, 27–41.

Suarez, J.A., Howard, J.D., Schoenbaum, G., Kahnt, T., 2019. Sensory prediction errors in the human midbrain signal identity violations independent of perceptual distance. Elife 8.

Sudlow, C., Gallacher, J., Allen, N., Beral, V., Burton, P., Danesh, J., Downey, P., Elliott, P., Green, J., Landray, M., Liu, B., Matthews, P., Ong, G., Pell, J., Silman, A., Young, A., Sprosen, T., Peakman, T., Collins, R., 2015. UK biobank: an open access resource for identifying the causes of a wide range of complex diseases of middle and old age. PLoS Med 12, e1001779.

Sutton, R.S., 1992. Gain adaptation beats least squares. Proceedings of the 7th Yale workshop on adaptive and learning systems.

Symmonds, M., Moran, C.H., Leite, M.I., Buckley, C., Irani, S.R., Stephan, K.E., Friston, K.J., Moran, R.J., 2018. Ion channels in EEG: isolating channel dysfunction in NMDA receptor antibody encephalitis. Brain 141, 1691–1702.

Takahashi, Y.K., Batchelor, H.M., Liu, B., Khanna, A., Morales, M., Schoenbaum, G., 2017. Dopamine Neurons Respond to Errors in the Prediction of Sensory Features of Expected Rewards. Neuron 95, 1395–1405 e1393.

Van Essen, D.C., Smith, S.M., Barch, D.M., Behrens, T.E.J., Yacoub, E., Ugurbil, K., Consortium, W.-M.H., 2013. The WU-Minn Human Connectome Project: An overview. Neuroimage 80, 62–79.

Van Horn, J.D., Grethe, J.S., Kostelec, P., Woodward, J.B., Aslam, J.A., Rus, D., Rockmore, D., Gazzaniga, M.S., 2001. The Functional Magnetic Resonance Imaging Data Center (fMRIDC): the challenges and rewards of large-scale databasing of neuroimaging studies. Philos Trans R Soc Lond B Biol Sci 356, 1323–1339.

Van Snellenberg, J.X., Torres, I.J., Thornton, A.E., 2006. Functional neuroimaging of working memory in schizophrenia: task performance as a moderating variable. Neuropsychology 20, 497–510.

Varoquaux, G., 2018. Cross-validation failure: Small sample sizes lead to large error bars. Neuroimage 180, 68–77.

Varoquaux, G., Raamana, P.R., Engemann, D.A., Hoyos-Idrobo, A., Schwartz, Y., Thirion, B., 2017. Assessing and tuning brain decoders: Cross-validation, caveats, and guidelines. Neuroimage 145, 166–179.

Verstynen, T.D., Deshpande, V., 2011. Using pulse oximetry to account for high and low frequency physiological artifacts in the BOLD signal. Neuroimage 55, 1633–1644.

Volz, S., Callaghan, M.F., Josephs, O., Weiskopf, N., 2019. Maximising BOLD sensitivity through automated EPI protocol optimisation. Neuroimage 189, 159–170.

Vossel, S., Bauer, M., Mathys, C., Adams, R.A., Dolan, R.J., Stephan, K.E., Friston, K.J., 2014. Cholinergic stimulation enhances Bayesian belief updating in the deployment of spatial attention. J Neurosci 34, 15735–15742.

Wang, L., Hermens, D.F., Hickie, I.B., Lagopoulos, J., 2012. A systematic review of resting-state functional-MRI studies in major depression. J Affect Disord 142, 6–12.

Wang, X.J., Krystal, J.H., 2014. Computational psychiatry. Neuron 84, 638–654.

Weber, L.A., Diaconescu, A.O., Mathys, C., Schmidt, A., Kometer, M., Vollenweider, F.X., Stephan, K.E., 2020. Ketamine Affects Prediction Errors About Statistical Regularities: A Computational Single-Trial Analysis of the Mismatch Negativity. J. Neurosci.

Weldon, K.B., Olman, C.A., 2021. Forging a path to mesoscopic imaging success with ultra-high field functional magnetic resonance imaging. Philos Trans R Soc Lond B Biol Sci 376, 20200040.

Wiecki, T.V., Sofer, I., Frank, M.J., 2013. HDDM: Hierarchical Bayesian estimation of the Drift-Diffusion Model in Python. Frontiers in Neuroinformatics 7.

Wipf, D., Nagarajan, S., 2009. A unified Bayesian framework for MEG/EEG source imaging. Neuroimage 44, 947–966.

Woodard, J.P., Carley-Spencer, M.P., 2006. No-reference image quality metrics for structural MRI. Neuroinformatics 4, 243–262.

Woolrich, M.W., Stephan, K.E., 2013. Biophysical network models and the human connectome. Neuroimage 80, 330–338.

World Health Organization, 1990. International Classification of Diseases. World Health Organization Press.

Yao, Y., Raman, S.S., Schiek, M., Leff, A., Frassle, S., Stephan, K.E., 2018. Variational Bayesian inversion for hierarchical unsupervised generative embedding (HUGE). Neuroimage 179, 604–619.

Yao, Y., Stephan, K.E., 2020. Markov chain Monte Carlo methods for hierarchical clustering of dynamic causal models. arXiv:2012.05744.

Yendiki, A., Koldewyn, K., Kakunoori, S., Kanwisher, N., Fischl, B., 2014. Spurious group differences due to head motion in a diffusion MRI study. Neuroimage 88, 79–90.

Yousefi, A., Paulk, A.C., Basu, I., Mirsky, J.L., Dougherty, D.D., Eskandar, E.N., Eden, U.T., Widge, A.S., 2018. COMPASS: An Open-Source, General-Purpose Software Toolkit for Computational Psychiatry. Front Neurosci 12, 957.

Yu, A.J., Dayan, P., 2005. Uncertainty, neuromodulation, and attention. Neuron 46, 681–692.

